# A benchmark of protein solubility prediction methods on UDP-dependent glycosyltransferases

**DOI:** 10.1101/2020.02.28.962894

**Authors:** Fatemeh Ashari Ghomi, Tiia Kittilä, Ditte Hededam Welner

**Affiliations:** Technical University of Denmark, Novo Nordisk Foundation Center for Biosustainability

**Keywords:** Protein solubility, Benchmark, UDP-dependent glycosyltransferase, Heterologous expression, *Escherichia coli*, SoluProt

## Abstract

UDP-dependent glycosyltransferases (UGTs) are enzymes that glycosylate a wide variety of natural products, thereby modifying their physico-chemical properties, i.e. solubility, stability, reactivity, and function. To successfully leverage the UGTs in biocatalytic processes, we need to be able to screen and characterise them *in vitro*, which requires efficient heterologous expression in amenable hosts, preferably *Escherichia coli*. However, many UGTs are insoluble when expressed in standard and attempted optimised *E. coli* conditions, resulting in many unproductive and costly experiments. To overcome this limitation, we have investigated the performance of 11 existing solubility predictors on a dataset of 57 UGTs expressed in *E. coli*. We show that SoluProt outperforms other methods in terms of both threshold-independent and threshold-dependent measures. Among the benchmarked methods, only SoluProt is significantly better than random predictors using both measures. Moreover, we show that SoluProt uses a threshold for separating soluble and insoluble proteins that is optimal for our dataset. Hence, we conclude that using SoluProt to select UGT sequences for *in vitro* investigation will significantly increase the success rate of soluble expression, thereby minimising cost and enabling efficient characterisation efforts for biocatalysis research.

## Introduction

Plants produce a wide range of natural product glycosides, which are used in industry for example as scents, dyes, flavours, and cosmetics [1–3]. Glycosylation can alter solubility, stability, reactivity and function of natural products and therefore glycosylation routes are of high importance [1]. In plants, natural products are glycosylated by UDP-dependent glycosyltransferases (UGTs) [4,5]. A single plant can have hundreds of UGTs, but unfortunately it is not well understood what governs the selectivity of these enzymes, limiting their usability in biotechnological applications. Their application is further hampered by the fact that many UGTs are poorly soluble when expressed in heterologous hosts, making screening, characterisation, and production efforts costly. For example, from our library of 57 UGT sequences of various plant origin expressed in *Escherichia coli*, only 31 sequences (54%) were soluble in the tested conditions.

*E. coli* is one of the most common host organisms for heterologous expression of recombinant proteins due to its versatile technological toolbox, availability, high growth rate, and continuous fermentation potential [6,7]. However, there is no guarantee for solubility of recombinant target proteins. For example, over-expression of recombinant proteins can exhaust the bacterial quality control system resulting in the formation of aggregates of misfolded and partially folded proteins called inclusion bodies [8]. In some cases, this can be exploited to simplify downstream processing [9], but in many cases, recovery from inclusion bodies is inefficient at best [10,11]. Nonetheless, *E.coli* remains the host of choice for recombinant protein expression in the academic research community in general as well as the UGT research community, since no other host has so far been identified as the silver bullet for recombinant UGT expression.

Solubility prediction software tools can have a significant impact on recombinant protein production by excluding insoluble proteins from expression trials and thereby preventing extra costs and dead-end experiments. Overall, solubility prediction tools can be grouped into 3 classes based on their applications [12]: 1) methods that predict the overall solubility of proteins upon expression (usually in *E. coli*), 2) approaches for predicting the aggregation propensity of different regions in a protein sequence, and 3) tools that predict the impact of mutations on solubility of proteins. Among these groups, the former is studied here.

Some of the existing protein solubility prediction tools have previously been benchmarked on a dataset of 2000 non-redundant proteins [13] from different families, which showed an accuracy range of 51 - 64 (%) for the five benchmarked methods (PROSOII, ccSOL, SOLpro, PROSO, and RPSP) with PROSOII being the most accurate predictor. In this study, we have benchmarked the performance of 11 protein solubility prediction tools on a dataset of 57 plant UGT proteins expressed in *E. coli* which were not used in other benchmarks. UGTs are notoriously difficult to handle recombinantly, and together with their high industrial relevance, this makes them an interesting target for *in silico* protein solubility prediction. We show here that SoluProt is the best predictor for UGT solubility, using both threshold-dependent and threshold-independent analyses. Moreover, this method uses a threshold that separates soluble and insoluble proteins well.

## Results

We have compared the performance of 11 sequence based solubility prediction methods on plant UGT proteins. We have only included the tools that predict the solubility of different proteins expressed in *E. coli* [12,13]. The list of the tools and their description can be found in the Methods section. These methods were tested on a dataset of plant UGT sequences. The dataset consists of 57 plant UGT sequences with their experimentally validated solubility (Methods), from which 31 sequences are soluble and 26 are insoluble. We have statistically analysed the results from different methods of solubility prediction for UGTs and compared them to experimentally validated data.

### Threshold-independent comparison

In order to compare the solubility prediction of different methods on UGT sequences, we used Receiver Operating Characteristic (ROC) curves shown in Figure 1A. For each method, the UGTs are sorted based on their solubility measure in increasing order. A ROC curve starts from point (0,0) and the first protein in the sorted list. If the protein is actually insoluble, it goes one step upwards and if it is soluble it moves one step to the right. In case the predictor works perfectly, the curve should first move *n* steps upwards followed by *m* steps to the right, where *n* and *m* represent the number of insoluble and soluble proteins, respectively. This results in an Area Under the Curve (AUC) of 1. However, if all of the predictions are incorrect, the AUC will be zero and a random predictor has an AUC of around 0.5. Overall, higher AUC values mean better predictions. The comparison shows that the AUC values for 4 of the methods are below 0.5 However, SoluProt (AUC = 66.4) followed by DeepSol3 (AUC = 65.3), and SOLpro (AUC = 63.2) are the three top performing tools (Figure 1B).

**Figure 1.**
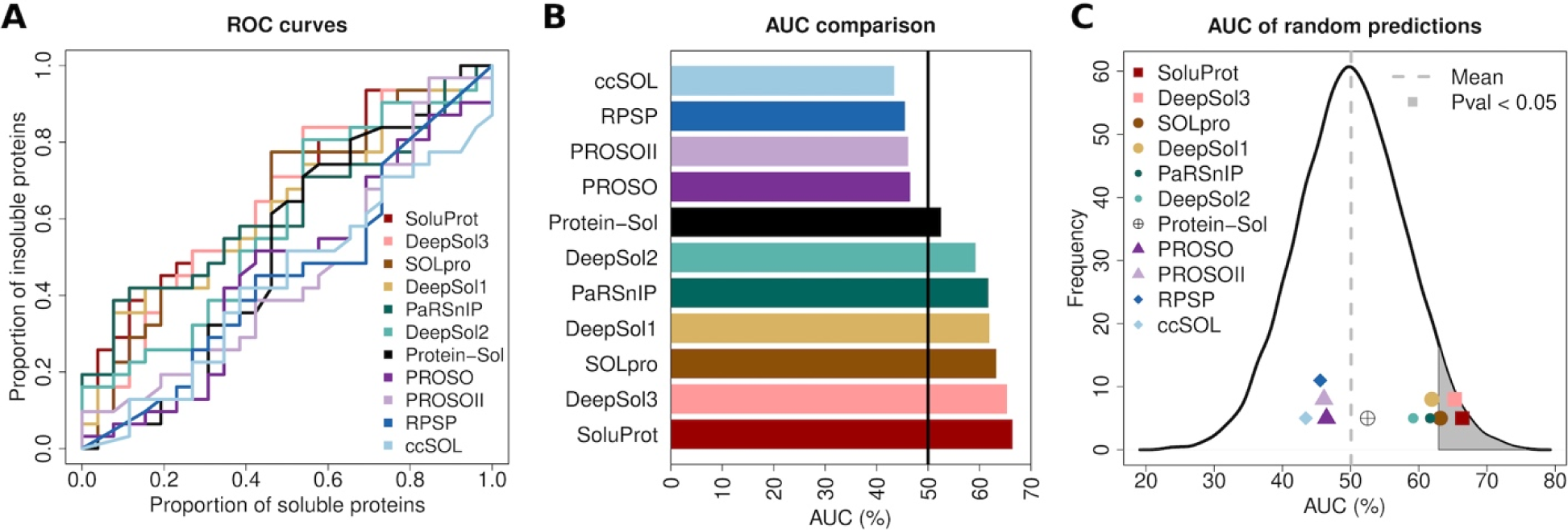
Threshold independent evaluation of the 11 solubility prediction methods on UDP-dependent Glycosyltransferase (UGT) dataset. A) Receiver Operating Characteristic (ROC) curves comparing the methods. The x axis shows the ratio of visited soluble proteins at each step to the whole number of soluble proteins and the y axis shows the ratio of visited insoluble proteins to the number of all insoluble proteins. B) Area under the ROC curves (AUC) comparing 11 different methods of solubility prediction shown in percentage. The higher the AUC, the better the method and a random method would have an AUC of about 50%, which is shown with a vertical line. C) The curve shows the distribution of AUC values obtained from 10,000 random solubility prediction methods. The dashed line shows the average AUC value obtained from simulated predictors which is about 0.5. The grey area on the right shows the top 5%, which corresponds to P-value < 0.05. SoluProt, DeepSol3, and SOLpro are the only predictors in this area.

Since AUC values were close to random for many of the methods, we then asked whether any of the predictors work significantly better than a random predictor for evaluating UGT solubility. To this end, we simulated 10,000 predictors, each of which randomly assigns a solubility score to each UGT protein using a uniform distribution. Afterwards, we calculated the AUC values for every single predictor and plotted their distribution together with the AUC values for the 11 benchmarked methods (Figure 1C). The top 5% of simulated predictors are shown in the grey shaded area which is equivalent to having less than 5% chance of obtaining these results by random i.e. P-value < 0.05. As the figure shows, the difference between the predictions from the benchmarked methods and random predictors is statistically significant for SoluProt, DeepSol3 and SOLpro.

### Threshold-dependent comparison

So far, we have used AUCs to evaluate the goodness of different methods. This measure is independent of the thresholds used to separate soluble and insoluble proteins. Nevertheless, this is not the only measurement for evaluating the performance of predictors. In Figure 2, we have used 3 other measures. Since these measures rely on a threshold for separating soluble and insoluble proteins, we excluded Protein-Sol from this analysis which does not provide a threshold. Sensitivity is the number of soluble UGTs that are correctly predicted as soluble divided by the number of all soluble UGTs, whereas specificity is the number of insoluble UGTs that are correctly predicted as insoluble divided by the number of all insoluble UGTs. Specificity is an important measure when one intends to select only soluble proteins for their study. Among all methods, SoluProt and SOLpro share the first place in terms of specificity (84.6). However, sensitivity and specificity very much depend on the number of soluble and insoluble predictions. For example, a method that predicts every protein as soluble has 100% sensitivity and 0% specificity, while a method that predicts everything as insoluble has 0% sensitivity and 100% specificity. It can be seen in Figure 2A that for most of the methods there is almost an inverse relationship between sensitivity and specificity. In order to take both sensitivity and specificity into consideration, we have used the balanced accuracy measure. Balanced accuracy is the average of sensitivity and specificity and detects the methods that work well based on both measures. In agreement with the threshold-independent analysis, the comparison of balanced accuracy for different methods shows that SoluProt (61.7), SOLpro (58.4), and DeepSol3 (57.3) outperform other methods (Figure 2B). We also compared the balanced accuracy of the prediction methods with that of 10,000 random predictors that randomly assign a solubility score to each sequence. A protein is considered soluble if its assigned score is above 0.5 and insoluble otherwise. We showed that SoluProt is the only method whose balanced accuracy is significantly higher than random predictors (Figure 2C).

**Figure 2.**
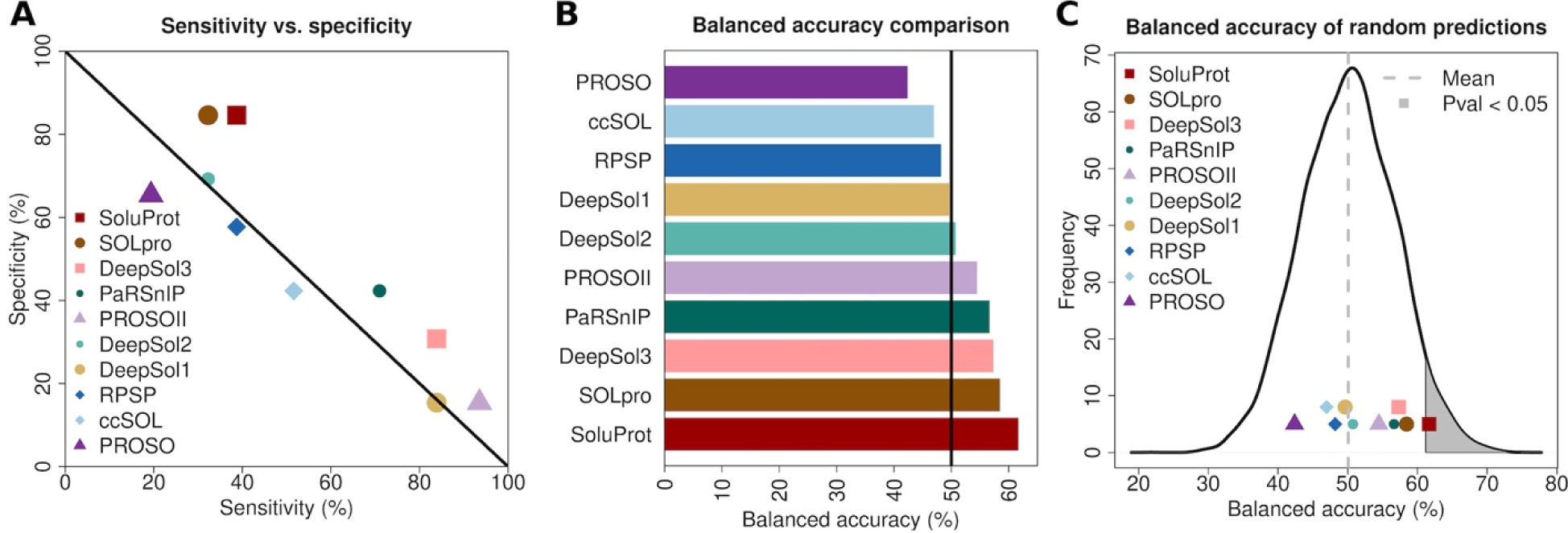
Threshold dependent evaluation of the solubility prediction methods. A) The comparison of sensitivity and specificity for all of the benchmarked methods for predicting soluble proteins on UGT dataset. Protein-Sol is excluded from this figure since it does not provide a threshold for separating soluble and insoluble proteins. The points around the black line have an inverse relationship between sensitivity and specificity. The best predictors are the ones above the line that are most distant from it. B) The comparison of balanced accuracy for the benchmarked methods (excluding Protein-Sol). The vertical line shows the average balanced accuracy of a random method. C) The curve shows the distribution of balanced accuracy values obtained from 10,000 random solubility prediction methods. The dashed line shows the average AUC value obtained from simulated predictors which is about 0.5. The grey area on the right shows the top 5%, which corresponds to P-value < 0.05. SoluProt is the only method in this region.

We also asked whether the proteins with highest solubility scores from each method are actually soluble and proteins with lowest solubility scores are insoluble. To this end, we sorted the UGTs based on their solubility scores for each method (Figure 3). As the figure shows, looking at the top 10% predictions (top 5) from each method, SoluProt, DeepSol3, PaRSnIP, and DeepSol2 predict no false positives. Moreover, for the top 20% predictions (top 11), SoluProt, DeepSol1, and PaRSnIP are the best predictors with 2 false positives. This indicates that using these methods with higher thresholds for separating soluble and insoluble proteins can result in very few false positives.

**Figure 3.**
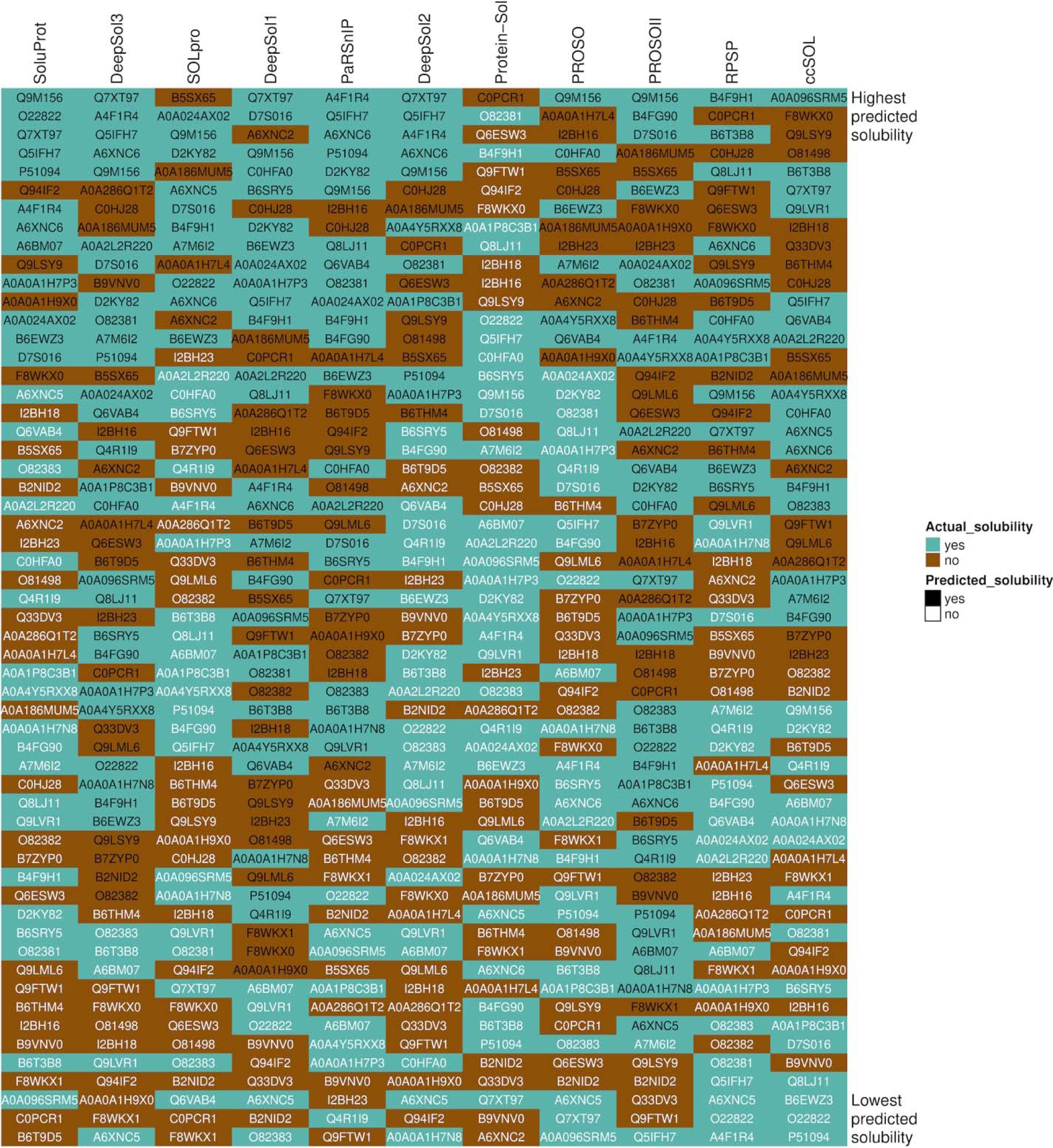
A comparison of solubility predictions using the 11 benchmarked methods. Each column shows the predictions of a method sorted by their solubility score so that the highest predicted solubility is at the top and the lowest predicted solubility is at the bottom. Brown shows experimental solubility and turquoise shows experimental insolubility. In all columns apart from Protein-Sol, protein names in black are predicted as soluble and proteins written in white are predicted as insoluble. For protein-Sol black shows a solubility score higher than average soluble *E. coli* proteins whereas white shows solubility scores lower than it.

### Threshold evaluation

In order to test whether the suggested thresholds for discriminating between soluble and insoluble UGTs are correctly assigned for different methods, we calculated the balanced accuracy of the predictions using different thresholds. These values are visualised in Figure 4. The dashed light brown lines show the thresholds defined by the methods and the solid turquoise lines show the best thresholds that maximise balanced accuracy for our dataset. As Protein-Sol does not provide a threshold for separating soluble and insoluble proteins, we have used a dotted dark brown line for it which visualises the average solubility score for soluble proteins in *E. coli*. The figures show that apart from SoluProt, DeepSol3, and PROSOII, in all the other methods, the predefined thresholds are distant from the best thresholds. PROSOII is the only method that uses a threshold of 60%, rather than 50%, for distinguishing between soluble and insoluble proteins and Figure 4 shows that this threshold actually yields the maximum possible accuracy for this method.

**Figure 4.**
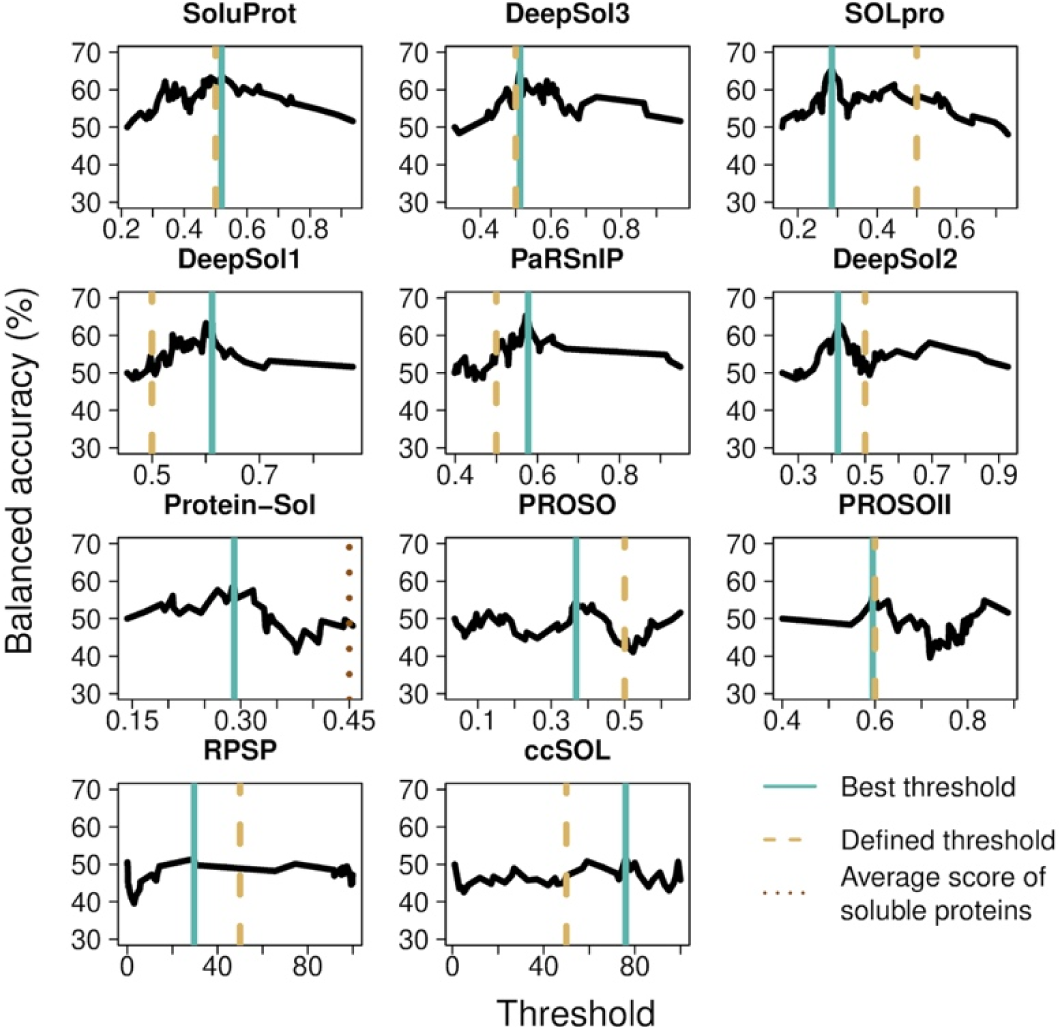
The comparison of the pre-defined thresholds for predicting solubility (dashed light brown) and the best thresholds that maximise balanced accuracy (solid turquoise). The dotted dark brown line for Protein-Sol shows the average solubility score of soluble *Escherichia coli* proteins. In all cases except for SoluProt, DeepSol3, and PROSOII, the defined thresholds are distant from the best thresholds, while in these three cases they are almost equal.

## Discussion

In this study, we compared different solubility prediction methods for predicting the solubility of UGT sequences when expressed in *E. coli*. For this, we used both threshold-dependent and threshold-independent measures. We showed that among all of the methods SoluProt is the best performer in terms of AUC, which is a threshold-independent measure, followed by DeepSol3, and SOLpro (Figure 1A and B). Moreover, these three are the only methods that outperform random predictors in a statistically significant manner (Figure 1C). For a threshold-dependent comparison, we calculated balanced accuracy values and found that the best tools are SoluProt, SOLpro, and DeepSol3 (Figure 2B). However, only SoluProt performs significantly better than random predictors (Figure 2C). We also investigated the solubility thresholds in different methods and showed that the solubility threshold for the majority of methods is distant from the best threshold that maximises their accuracy. This analysis shows that proposed thresholds for most of the methods are not optimal for UGT proteins. The exceptions to this are SoluProt, DeepSol3, and PROSOII (Figure 5).

Chang et al. [13] have previously reported accuracy values comparing PROSOII (64.35), SOLpro (59.95), PROSO (57.85), ccSOL (54.20), and RPSP (51.45). The ranking of the tools based on accuracy from Chang et al. [13] is to some extent different from our study (SOLpro, PROSOII, RPSP, ccSOL, and PROSO). In addition, the accuracy values in Chang et al. [13] are generally higher than our findings. One reason for these differences is that the test data in Chang et al. [13] is randomly selected from the datasets used for the development of these tools which results in higher accuracy for the tools and also works in favour of the tools with larger datasets. As a matter of fact, the reported accuracy values in the mentioned benchmark and the size of datasets are highly correlated with 0.88 Pearson’s correlation coefficient. Other factors for the dissimilarities in the results are the small size of our dataset compared to their dataset of 2000 proteins and the differences in the nature of the datasets since their dataset is diverse, while ours is homogeneous and only includes plant UGTs.

Overall, this analysis shows that SoluProt outperforms all the other methods for predicting the solubility of plant UGTs and its predictions are significantly better than random predictors using threshold-dependent and threshold-independent measures. This method can be used for selecting soluble enzymes for *in vitro* studies.

## Methods

### Solubility dataset

We used a set of 57 plant UGT sequences with 31 soluble and 26 insoluble proteins. The whole dataset with UniProt IDs and their solubility status as well as their predicted solubility using the benchmarked methods is provided in Supplementary Materials. The sequences in fasta format can also be found in Supplementary Materials. In this file, 5 sequences are annotated with the word “indigo” in their header. The solubility of these sequences was studied according to Hsu et al. [14]. In addition, we added all of the plant based non-redundant BLAST hits of the protein with UniProt ID B6SRY5 in PDB database to the soluble group. We assumed these proteins are soluble, since they have a solved structure on PDB. These are annotated with “BLAST”. The solubility of the rest of the sequences are studied as explained in the following section.

### Experimental solubility identification

Synthetic genes encoding different UGTs were obtained from Genscript in a modified pET28a(+) vector with an N-terminal His-tag followed by a TEV-cleavage site and the gene of interest. Plasmids were transformed into BL21 Star (DE3) cells (Invitrogen) for expression. 80 mL of Luria-Bertani media was supplemented with kanamycin (50 µg/mL), inoculated with 1 % overnight culture and grown at 37°C until OD_600_ reached 0.5-0.8. Protein expression was induced with 0.5 mM IPTG and cells grown overnight at 18°C. Cells were harvested with centrifugation (4000xg, 15 min, 4°C) and stored at −20°C.

All purification steps were done on ice or in a cold room. Cell pellets were thawed and dissolved in lysis buffer (50 mM Hepes, 300 mM NaCl, 20 mM Imidazole, 1 mM DTT, pH 7.4 supplemented with 1 µg/mL Dnase I and cOmplete EDTA-free protease inhibitor cocktail (Roche)). Cells were lysed via three passes through EmulsiFlex C5 (Avestin) and the lysate was cleared with centrifugation (12,000xg, 40 min, 4°C). Supernatant was incubated with Ni-NTA beads (HisPur NiNTA resin, Thermo-Fischer) with gentle shaking (1h) and the beads were washed three times with wash buffer (50 mM Hepes, 300 mM NaCl, 20 mM Imidazole, pH 7.4). Bound proteins were eluted with elution buffer (50 mM Hepes, 300 mM NaCl, 250 mM Imidazole, pH 7.4). Samples of different purification steps were analysed on SDS-PAGE (NuPAGE 4-12 % Bis-Tris protein gel) to assess solubility of proteins.

### Solubility prediction tools

Here we will describe how we ran each of the methods and whether we have performed any processing on the output of the tools to prepare them for the statistical analyses. The predictors are selected from a review paper by Musil et al. [12] and a benchmark by Chang et al. [13]. We have excluded some of the tools which we could not access [15] or that predicted solubility of proteins expressed in the periplasm of *E. coli* which is not tested in this experiment [16]. All of the statistical analyses described in this paper are provided as an R script on https://github.com/ftmashari/Solubility_prediction.

**SoluProt** [17] uses 36 sequence-based features and predicts solubility score of proteins using random forest regression, given that they are expressed in *E. coli*. The tool can be run from https://loschmidt.chemi.muni.cz/soluprot/.

**DeepSol** [18] uses a deep learning approach called Convolutional Neural Network and feeds it with protein sequences as well as sequence based features extracted from them and secondary structure based features calculated using SCRATCH. DeepSol has 3 different configurations that differ in their network inputs. The output of the program is a score for solubility of proteins given that they are expressed in *E. coli*. DeepSol was downloaded from https://github.com/sameerkhurana10/DSOL_rv0.2 and installed and run according to the guidelines.

**SOLpro** [19] uses an ensemble of 20 Support Vector Machine (SVM) classifiers that are trained based on different features extracted from protein sequences as well as secondary structure features predicted using SCRATCH suite [20]. It then feeds the output of these classifiers in addition to the normalised sequence lengths to another SVM classifier. The output of this SVM classifier shows the solubility propensity of proteins on expression in *E. coli*. SOLpro was used through its website (http://scratch.proteomics.ics.uci.edu) by providing the sequence and selecting the SOLpro button. For the proteins that were predicted as insoluble, we subtracted the insolubility score from 1 to obtain the solubility score.

**PaRSnIP** [21] uses a set of features extracted from protein sequences in addition to another set of secondary structure features predicted using SCRATCH suite. It then uses Gradient Boosting Machines to provide a score for solubility upon expression in *E. coli*. PARSnIP was downloaded from https://github.com/RedaRawi/PaRSnIP and installed and run according to the guidelines.

**Protein-Sol** [22] extracts 35 sequence based properties from proteins and using these features calculates a measure for solubility. This measure is then compared to values obtained from *E. coli* proteins in a cell-free expression system [23]. Protein-Sol was run from https://protein-sol.manchester.ac.uk and the predicted scaled solubility value was used for this analysis.

**PROSO** [24] is an older version of PROSO II which uses a Naive Bayes classifier to combine the output of SVM classifiers and provide a score for solubility of proteins expressed in *E. coli*. PROSO was run from http://mbiljj45.bio.med.uni-muenchen.de:8888/proso/proso.seam.

**PROSO II** [25] extracts a set of sequence based and global features from proteins. First, it uses a parzen window classifier in addition to logistic regression for predicting solubility. It then uses another logistic regression algorithm to combine the results obtained from the first stage and provides a solubility score upon heterologous expression in *E. coli*. PROSO II was run from http://mbiljj45.bio.med.uni-muenchen.de:8888/prosoII/prosoII.seam by providing sequences in fasta format. Contrary to other methods that use a threshold of 0.5 for solubility score, in this method sequences with scores greater than or equal to 0.6 are considered soluble.

**RPSP** [26,27] extracts 32 features from protein sequences and using logistic regression predicts a score for proteins to be soluble given that they are expressed in *E. coli*. RPSP was run using its web page (http://biotech.ou.edu). The average pI and molecular weight were calculated using https://web.expasy.org/compute_pi/.

**ccSOL omics** [28] feeds the physico-chemical properties of proteins to a neural network with one hidden layer to predict the solubility propensity of proteins upon expression in *E. coli*. ccSOL omics was run from http://s.tartaglialab.com/update_submission/227788/66268c5911.

### Key points

1. Among the 11 methods benchmarked for predicting the solubility of UDP-Glycosyltransferases, 8 fail to perform significantly better than random predictors both in threshold-dependent and threshold-independent comparisons.
2. SOLpro and DeepSol3 are significantly better than random predictors using threshold-independent comparisons, but fail to perform well in threshold-dependent comparisons.
3. SoluProt is the only method that performs better than random in both comparisons while it also outperforms all the other tools.

## Acknowledgements

The authors would like to thank Daniel Dunn for technical assistance.

## Author contributions

FA has designed the benchmarking analyses, tested all of the solubility prediction methods, performed the statistical analyses, drafted and revised the manuscript. TK has designed and performed the laboratory experiments, evaluated the solubility of the UGT dataset, drafted and revised the manuscript. DHW has designed the experiments and analyses, evaluated the solubility of UGT dataset, supervised both the experimental and analytical sides of the project, provided funding, and revised the manuscript. All of the authors have read the article and approved it.

