## Supplementary Materials for "A benchmark of protein solubility prediction methods on UDP-dependent glycosyltransferases"

| UGT | Experimental | SOLpro | ccSol_omic | RPSP | PaRSnIP | PROSOII | PROSO | Protein_Sol | DeepSol1 | DeepSol2 | DeepSol3 | SoluProt |
| --- | --- | --- | --- | --- | --- | --- | --- | --- | --- | --- | --- | --- |
| B6SRY5 | yes | 0.443206 | 4 | 74.5 | 0.5399 | 0.685 | 0.191 | 0.356 | 0.646413 | 0.49658957 | 0.56789315 | 0.3433 |
| B4F9H1 | yes | 0.578745 | 77 | 100 | 0.5778 | 0.713 | 0.161 | 0.412 | 0.611953 | 0.47662717 | 0.51322645 | 0.3557 |
| B6THM4 | no | 0.268868 | 97 | 91.8 | 0.4695 | 0.796 | 0.397 | 0.252 | 0.569969 | 0.5004781 | 0.490186 | 0.3138 |
| B6T9D5 | no | 0.267455 | 42 | 99.2 | 0.5622 | 0.691 | 0.356 | 0.286 | 0.572549 | 0.4901107 | 0.57816344 | 0.2188 |
| A0A096SRM5 | yes | 0.258425 | 100 | 99.6 | 0.4451 | 0.729 | 0.037 | 0.333 | 0.561406 | 0.4186741 | 0.5775597 | 0.2614 |
| B7ZYP0 | no | 0.406081 | 57 | 3.8 | 0.5347 | 0.759 | 0.357 | 0.259 | 0.538093 | 0.471811 | 0.5056163 | 0.3634 |
| C0PCR1 | no | 0.163681 | 17 | 100 | 0.5368 | 0.719 | 0.088 | 0.455 | 0.606348 | 0.55821973 | 0.549077 | 0.237 |
| C0HFA0 | yes | 0.443992 | 82 | 98.9 | 0.5523 | 0.76 | 0.572 | 0.362 | 0.655504 | 0.30612385 | 0.58879405 | 0.4569 |
| B4FG90 | yes | 0.29037 | 59 | 0.4 | 0.5775 | 0.862 | 0.371 | 0.207 | 0.567075 | 0.49196666 | 0.56641334 | 0.4003 |
| B6T3B8 | yes | 0.324703 | 99 | 100 | 0.5255 | 0.716 | 0.1 | 0.206 | 0.548216 | 0.46690756 | 0.48395413 | 0.2896 |
| C0HJ28 | no | 0.26086 | 95 | 100 | 0.6051 | 0.798 | 0.564 | 0.334 | 0.634729 | 0.66726667 | 0.682687 | 0.3762 |
| D7S016 | yes | 0.583925 | 3 | 10.9 | 0.5402 | 0.836 | 0.411 | 0.342 | 0.718576 | 0.4834545 | 0.6523582 | 0.5188 |
| Q4R1I9 | yes | 0.391572 | 37 | 1.1 | 0.4027 | 0.676 | 0.416 | 0.292 | 0.521804 | 0.478163 | 0.59322727 | 0.4528 |
| B2NID2 | no | 0.198657 | 46 | 96.2 | 0.4478 | 0.568 | 0.064 | 0.201 | 0.465008 | 0.45154685 | 0.50142145 | 0.4625 |
| Q8LJ11 | yes | 0.323394 | 1 | 100 | 0.6036 | 0.631 | 0.457 | 0.388 | 0.600425 | 0.43291602 | 0.574233 | 0.3757 |
| Q9FTW1 | no | 0.4105 | 74 | 100 | 0.3987 | 0.548 | 0.158 | 0.41 | 0.559155 | 0.31583324 | 0.46769008 | 0.3221 |
| Q6ESW3 | no | 0.23518 | 33 | 100 | 0.4897 | 0.787 | 0.076 | 0.442 | 0.59406 | 0.53826416 | 0.5781873 | 0.3514 |
| Q94IF2 | no | 0.251877 | 7 | 95 | 0.5613 | 0.788 | 0.29 | 0.409 | 0.490853 | 0.2933309 | 0.41685176 | 0.7282 |
| A6XNC2 | no | 0.54421 | 78 | 13.4 | 0.5045 | 0.777 | 0.523 | 0.143 | 0.708485 | 0.48630795 | 0.59313035 | 0.4622 |
| B5SX65 | no | 0.729899 | 88 | 5.9 | 0.4437 | 0.818 | 0.57 | 0.336 | 0.565505 | 0.5130822 | 0.61464286 | 0.4779 |
| I2BH23 | no | 0.490065 | 50 | 0 | 0.4065 | 0.804 | 0.551 | 0.314 | 0.534634 | 0.47426125 | 0.57152003 | 0.4594 |
| I2BH18 | no | 0.25647 | 98 | 14.3 | 0.5286 | 0.729 | 0.32 | 0.377 | 0.546581 | 0.3593473 | 0.430891 | 0.4822 |
| I2BH16 | no | 0.273984 | 4 | 0 | 0.6092 | 0.759 | 0.589 | 0.377 | 0.595779 | 0.41743845 | 0.60047233 | 0.3085 |
| A0A286Q1T2 | no | 0.350842 | 73 | 0 | 0.4306 | 0.738 | 0.529 | 0.293 | 0.597457 | 0.35563502 | 0.6894387 | 0.4198 |
| A0A186MUM5 | no | 0.6395 | 86 | 0 | 0.4967 | 0.833 | 0.561 | 0.259 | 0.610493 | 0.65088904 | 0.67879987 | 0.407 |
| B6EWZ3 | yes | 0.5 | 1 | 91.6 | 0.5714 | 0.808 | 0.563 | 0.288 | 0.620885 | 0.47334322 | 0.5121156 | 0.5467 |
| A6BM07 | yes | 0.306977 | 33 | 0 | 0.4286 | 0.633 | 0.317 | 0.334 | 0.499819 | 0.39051992 | 0.4745231 | 0.6375 |
| F8WKX1 | no | 0.160749 | 23 | 0 | 0.4675 | 0.607 | 0.167 | 0.245 | 0.511368 | 0.40824348 | 0.3389717 | 0.2796 |
| F8WKX0 | no | 0.237817 | 100 | 99.9 | 0.5693 | 0.808 | 0.229 | 0.407 | 0.510953 | 0.40299228 | 0.44941163 | 0.5111 |
| D2KY82 | yes | 0.641034 | 45 | 0.7 | 0.6385 | 0.769 | 0.459 | 0.332 | 0.6313 | 0.46988106 | 0.63124734 | 0.3498 |
| A0A0A1H7N8 | yes | 0.257268 | 30 | 29.6 | 0.5178 | 0.628 | 0.233 | 0.268 | 0.528315 | 0.25209105 | 0.51394814 | 0.402 |
| A0A0A1H7P3 | yes | 0.34364 | 72 | 0 | 0.4236 | 0.731 | 0.442 | 0.333 | 0.612487 | 0.50273883 | 0.53198534 | 0.5885 |
| A0A0A1H7L4 | no | 0.560928 | 26 | 0.5 | 0.572 | 0.758 | 0.628 | 0.215 | 0.592962 | 0.3962123 | 0.5832507 | 0.4179 |
| A0A0A1H9X0 | no | 0.261399 | 5 | 0 | 0.5307 | 0.805 | 0.505 | 0.288 | 0.505868 | 0.30163902 | 0.37726367 | 0.5724 |
| A7M6I2 | yes | 0.568085 | 66 | 1.8 | 0.4916 | 0.595 | 0.534 | 0.338 | 0.571898 | 0.4373987 | 0.6160826 | 0.3959 |
| A0A1P8C3B1 | yes | 0.306422 | 4 | 97.3 | 0.4417 | 0.712 | 0.096 | 0.399 | 0.55407 | 0.5294313 | 0.5926412 | 0.4178 |
| B9VNV0 | no | 0.383322 | 3 | 5.6 | 0.4087 | 0.649 | 0.107 | 0.169 | 0.494955 | 0.47291532 | 0.6404192 | 0.2897 |
| O82383 | yes | 0.211125 | 76 | 0 | 0.5283 | 0.717 | 0.087 | 0.314 | 0.453819 | 0.44571763 | 0.4841308 | 0.4738 |
| Q9LML6 | no | 0.32829 | 74 | 65.5 | 0.5412 | 0.788 | 0.371 | 0.28 | 0.526706 | 0.3691521 | 0.51888645 | 0.3343 |
| O82382 | no | 0.325495 | 49 | 0 | 0.5301 | 0.655 | 0.262 | 0.337 | 0.548636 | 0.40796342 | 0.5006308 | 0.3642 |
| O82381 | yes | 0.252212 | 12 | 0 | 0.5871 | 0.8 | 0.459 | 0.444 | 0.553013 | 0.54795283 | 0.6270093 | 0.3396 |
| Q9LSY9 | no | 0.261456 | 100 | 99.8 | 0.5575 | 0.592 | 0.092 | 0.377 | 0.53742 | 0.52871066 | 0.50790745 | 0.619 |
| Q33DV3 | no | 0.339488 | 98 | 12.8 | 0.4981 | 0.568 | 0.349 | 0.195 | 0.47325 | 0.34434348 | 0.52238935 | 0.4311 |
| A0A4Y5RXX8 | yes | 0.305839 | 84 | 98.5 | 0.4256 | 0.792 | 0.508 | 0.319 | 0.541912 | 0.594791 | 0.5266851 | 0.4172 |
| Q5IFH7 | yes | 0.286166 | 93 | 0 | 0.9217 | 0.4 | 0.386 | 0.371 | 0.612348 | 0.8624973 | 0.86832535 | 0.7464 |
| A6XNC5 | yes | 0.600852 | 81 | 0 | 0.4474 | 0.603 | 0.061 | 0.259 | 0.46974 | 0.29483873 | 0.32655394 | 0.4846 |
| O22822 | yes | 0.560373 | 1 | 0 | 0.4507 | 0.716 | 0.368 | 0.376 | 0.497222 | 0.4491924 | 0.51688874 | 0.8899 |
| P51094 | yes | 0.303593 | 1 | 0.5 | 0.6681 | 0.644 | 0.13 | 0.206 | 0.523086 | 0.5057004 | 0.61537546 | 0.7394 |
| Q6VAB4 | yes | 0.19516 | 92 | 0.3 | 0.5955 | 0.771 | 0.506 | 0.277 | 0.53831 | 0.4862481 | 0.6049674 | 0.4795 |
| Q7XT97 | yes | 0.250689 | 99 | 94.7 | 0.5365 | 0.74 | 0.043 | 0.19 | 0.873749 | 0.92729396 | 0.9681539 | 0.8237 |
| A6XNC6 | yes | 0.552427 | 80 | 99.9 | 0.913 | 0.7 | 0.191 | 0.228 | 0.576978 | 0.75954354 | 0.86464787 | 0.6426 |
| A4F1R4 | yes | 0.351922 | 18 | 0 | 0.9483 | 0.795 | 0.201 | 0.317 | 0.590563 | 0.83697015 | 0.8742906 | 0.7071 |
| O81498 | no | 0.213221 | 100 | 3.1 | 0.544 | 0.724 | 0.116 | 0.339 | 0.532096 | 0.52056056 | 0.44356316 | 0.456 |
| Q9M156 | yes | 0.697091 | 46 | 95.5 | 0.6366 | 0.887 | 0.652 | 0.346 | 0.672863 | 0.692302 | 0.7293033 | 0.9365 |
| A0A024AX02 | yes | 0.717049 | 27 | 0.3 | 0.5802 | 0.804 | 0.476 | 0.291 | 0.614544 | 0.40703776 | 0.60677063 | 0.5631 |
| A0A2L2R220 | yes | 0.462159 | 91 | 0.1 | 0.5418 | 0.78 | 0.175 | 0.334 | 0.602611 | 0.46504867 | 0.66397434 | 0.4624 |
| Q9LVR1 | yes | 0.256052 | 99 | 30.2 | 0.5142 | 0.64 | 0.14 | 0.317 | 0.498651 | 0.39470226 | 0.42229182 | 0.371 |

Table containing the experimental solubility evaluation of all proteins as well as the predictions using benchmarked methods.

All sequences used for the benchmark are provided in the following in fasta format. The solubility of the sequences annotated with “indigo” was studied according to Hsu et al. [[14]](https://paperpile.com/c/jlfTzB/rWDUi).  In addition, we added all of the plant based non-redundant BLAST hits of the protein with UniProt ID B6SRY5 in PDB database to the soluble group. We assumed these proteins are soluble, since they have a solved structure on PDB. These are annotated with “BLAST”. The solubility of the rest of the sequences is evaluated as described in Methods section.

>B6SRY5

MENANASSCRNGDGTQTPPPHVAMLATPGMGHLIPLAELAKRLAQRHGVTSTLITFASTASATQRAFLASMPPAVASMALPPVDMSDLPRDAAIETLMSEECVRAVPALTEALLSLKQRPTTTGRLVAFVTDLFGADAFDAARAAGVQRRYLFFPTNLTALTLMLHLPELDASIPGEFRDLAEPLRLPGCVPLPGTETMKPLQDKSNPSYRWMVHHGAKFREATAILVNSFDAVEPGPAEVLRQPEPGRPPVRTIGPLVRAEDGGGSKDDAPCPCVEWLDRQPAKSVIFVSFGSGGTLPAEEMRELALGLELSGQRFLWVVRSPSEGGVGNDNYYDSASKKDPFSYLPQGFLERTKDVGLVVPSWAPQPKVLAHQSTGGFLTHCGWNSTLESLVHGVPMLAWPLFADQRQNAVLLCDGVGAALRVPGAKGREDIAAVVRELMTAEGKGAAVRAKVEELQKAAAEGLRDGGATAAALAEVVKEWTSADDSVY

>B4F9H1

MKAGPPHVAMLATPGMGHLIPLAELAKRLAARRGATATLITFASAASATQRAFLASLPPSVAARALPPVDLSDLPRDAAIETLMTAECARSLPAIAAVLAELGETARLVAFVVDQFGMEAFNAAGVRAARCLFMPMNLHALSLVLHLPELAASVPGEFRDLAEPVRLPGCVPIPGPDIISPLQDRSNPSYAVMVNLAVRCREAAAAILVNSFDAVEPEAAEALRHPAEPGWPPVYPVGPLILQSESGGTGADVDGTPPRAACLEWLDRQPARSVVYVSFGSGGALPKEQMHELALGLERSGQRFLWVVRSPSDDEGTLNGNYYDAESKKDPFAYLPEGFVGRTKEVGLLVPSWAPQTQVLAHGATGGFLTHCGWNSTLESLVHGVPMVAWPLFAEQRLNAVMLSEGAGAAIRLPETKDKESIAAVVRELVEGEGKGAMVRAKVAQLQKAAAEGLREGGAATTALDEVMDKWEAEAN

>B6THM4

MAATPTIVLVPVWGIGHFVPMLEVGKRLLARSALPLTVTVLVMPLPAEAKRASEITEHIRQQEASGLAIRFHHLPAVEPPTDHSGIEEYISRYVQLYSPHVKAAVAGLTCPVAGVVVDIFCTALFDAAHQLGVPAYVYLITSAAMCALLLRSPTLDEEVAVEVEFEQMEGGVDVPGLPPVPASCLPTGLENRKIPTYRWFLYNGRRYMEAAGIIVNTIAEAEPRVLAAIADGRCTRGVPAPPVYSIGPVIPSTPPAEQQAQECVRWLDSQPPSSVVFLCFGSGGCFTAPQAHEIAHGLDRSGHRFLWVLRGTPEPGTKLPSDGNLAELLPADFLARTKDRGLVWPTKAPQKEILAHAAVGGFVTHCGWNSVLESLWHGVPMVPWPLGAEQHYNAFTLVADMGVAVALNVERKRKNFVEATELERAVKALMCDGETARKVRDKVMEIKAACRKAMEEGGSSNMSLQRLCDALVEGAVHPGK

>B6T9D5

MAATAYPTVVLIPLCVPGHLASMIEAGKRLLGRSRCPMSLTVLITQMTMSANLMSDVDDIMRREADSGLDIRFVHLPAVELPTVHHGLEDFMMRFIQLHATHVKEAVSGMSSPVAAVVVDYFCTTLFDVARELALPAYAYMPSGASMVALMLRLPALDGEVSGDFEAMEGTVDLPGMPPVPARLMPSPLMRKDPNFAWLVYHGNRFMEADGVIVNTVAELEPSILAAIADGLCVSRRRAPAVYPIGPVLPLKPPSAPGDGEQVVAQRHECVRWLDAQPPASVVLLCFGSMGGSFPSPQVREIADGLERSGHRFLWVLRGPPPPDGSKYPTDANVHELLPEGFLERTKGRGLVWPTWAPQKDILANPAVGGFVTHCGWNSILESLWHGVPMVPLPQFAEQHLNAFELVSVMGVAVAMQVDRKRGNFVEAAELERAVRCLMGGSEEEGRKAREKATEAKALSQNGVASGGSSDASVQKLAREILHKHDDKGCATASGESMGSVVVPASPARII

>A0A096SRM5

MAANGGDHTSARPHVVLLPSAGMGHLVPFARLAVALSEGHGCNVSVAAVQPTVSSAESRLLDALFVAAAPAVRRLDFRLAPFDESEFPGADPFFLRFEATRRSAPLLGPLLDAAEASALVTDIVLASVALPVARERGVPCYVLFTSSAAMLSLCAYFPAYLDAHAAAGSVGVGVGNVDIPGVFRIPKSSVPQALHDPDHLFTQQFVANGRCLVACDGILVNTFDAFEPDAVTALRQGSITVSGGFPPVFTVGPMLPVRFQAEETADYMRWLSAQPPRSVVYVSFGSRKAIPRDQLRELAAGLEASGKRFLWVVKSTIVDRDDTADLGGLLGDGFLERVQGRAFVTMGWVEQEEILQHGSVGLFISHCGWNSLTEAAAFGVPVLAWPRFGDQRVNAALVARSGLGAWEEGWTWDGEEGLTTRKEVAKKIKGMMGYDAVAEKAAKVGDAAAAAIAKCGTSYQSLEEFVQRCRDAERK

>B7ZYP0

MQAIKKTIVLYPGLFVSHFVPMMQLADVLLEEGYAVAVALIDLTMDQDVTLAAAVDRVASAKPSVTIHRLPRIQNPPAITDCGGDGLLWYFKTVKRYNDPLREFLCSLQQQQPARSVVHAVILDGPSADALDVTKELGIPAYTFYATNASAVAVFLQLPWTHAEGQPSFKELGDTPLSISGVPPMPASYLMPPMLDDPASETYKTMMRVSRRNPEPEGILVNTFASLEGRVLRALRDPLFLPIGDDGCRRMPPVYCVGPLVVGAGDGDGVGVGEAKEKHECLAWLDEQPERSVVFLCFGSLGAAAHSEEQLKEIAVGLERSGHRFLWVVRAPLPTEGVDPGRLFDPRADFDLCALLPAGFLERTRARGLVVKLWAPQVNVLNHRATGAFVTHCGWNSVMEAVTAGVPMLCWPMYAEQKMNSVVMVEEAGIGVDLVGWQQGLVNAEEVERKVKMVMEFKEGEQLRARVTAHRDAAAVAWKDGGSSRAAFGLFLSDVDNHVHKAKTWSSANGRDGKLGQS

>C0PCR1

MASLATSTPAAAAGPRPRPDRPHVVLVPSPGVGHLMPMAELARRLVSHHALAATLVTFNLSGDPDAKSAAVLSSLRAANVSTATLPAVPLDDLPDDASIETVLFEVIGRSIPHLRAFLRDVGSTAGAPLAALVPDFFATAALPLASELGVPAYIFFPSNLSALSVMRSAVELHDGAGAGEYRDLPDPLPLPGGVSLRREDLPSGFRDSKESTYAQLIDAGRQYRTAAGILANAFYEMDPATVEEFKKAAEQGRFPPAYPVGPFVRSSSDEGSVSSPCIEWLDLQPTGSVVYVSFGSAGTLSVEQTAELAAGLENSGHRFLWIVRMSSLNGEHSDDMGRNYCDGGDENDPLAWLPEGFLERTRGRGLAVSSWAPQVRVLSHPATAAFVSHCGWNSTLESISSGVPMVAWPLFAEQRVNAVDLSEKVGVALRLGVRPDDGLVGREEIAAVVRELMEGEDGRAVRRRTGDLQQAADLAWASDGSSRRALEEVVSRWKAGAIRRAAV

>C0HFA0

MEATAAETGLAQRPVVLYPSPGMGHLVSMIELGKLLGARGLPVTIVVVEPPFNTGATAPFLAGVSAANPSISFHRLPKVERLPLVSTKHQEALTFEVIRVSNPHLREFLAAATPAVLVVDFFCSIALDVAEELRVPAYFFFTSGAEVLAFFLHLPALHERATASFQDMGEEPVQVPGIPPFPATHAILPVMERDDAAYDGFVKGCADLCRSQGVLVNTFRLLEQRAVETVAAGHCTPPGLPTPPIYCIGPLIKSEEVLGKGGEECLAWLDAQPRASVVLLCFGSIGRFSAEQIREVAAGLEASRQRFLWVVRAPPSDDPAKKFEKPPEPDLDALLPEGFLARTKDRGLVVKSWAPQRDVLAHASVGGFVTHCGWNSVLEAIMAGVPMVAWPLYAEQRLNRVFLEKEMQLAVAVAGYDSDKGLVPAEEVAAKVRWIMDSEGGRMLRERTLAAMRQAKDALREGGESEATLAGLVDDWKRT

>B4FG90

MKQQQTVILYPSPGVGHIVPMVQLAKVFLRHGCDVTMVIAEPAASSPDFRIVDLDRVAASNPAITFHVLPPVPYADLAVPGKHHFLLTLQVLRRYNGELERFLRSVPRERLHSLVVGMFCTDAVDVGAKLGVPVYTFFASAAATLAVVAQLPALLSGRRAGLKELGDTPLQFLGVPPFPASHLVRELLEHPDDDELCKTMVDVWKRCTDGSGVLVNTFESLESPAVQALRDPRCVPGRVLPPVYCVGPLIGGDGGTRAAAEQERAAETRHECLAWLDEQPENSVVFLCFGSRCAHSAEQLRGIAVGLERSGQRFLWSVRTPAGTDGGSENLGALFPEGFLQRTKDRGLVVRSWAPQVEVLRHPSTGAFMTHCGWNSTLEAITAGVPMLCWPFYAEQLMNKVFVTEGMGVGVEMEGYTTGFIKSEEVEAKVRLVMESEEGRHLRGRAVALKNEAQAALRDDGPSETSFARFLFDAKNLQGHLGHGSE

>B6T3B8

MAAPTVVLLPVWGAGHLMSMLDAGKRLLTRGGRALSLTVLVMRAPTEQLAADLDAHIRREEASGLDVRFVRLPAVQPPTHFHGIEEFISRLVQLHAPHVRAAISSLASPVAAVVMDFFCTALLDVTRELAVPAYVYFTASAGMLAFFLRLPSLHEEVTVQFEEMEGAVDVPGLPPVPPSSLPVPVMDKNHPNYTWFMYHGRRFAEADGIIVNTAAELEQSVLAAIADGRCTPGVRAPTVYPIGPVISFSPPPTNTEHPHECVRWLDTQPAASVVLLCFGSQGFSAAPQAHEIAHGLERSGHRFLWVLRGPPAPGERHPSDANLSELLPDGFLERTKGRGLVWPTKAPQKEILAHAAVGGFVTHGGWNSVLESLWFGVPMAPWPLYAEQHLNAFTLVAYVGVAVAMKVDRKRNNFVEASELERAVKELMGGGEEGRKAREKAMEMRDACRNAVEEGGSSYSSLRRLSEKICKVDKNL

>C0HJ28

MAHKTVILYPSLGVGHLNPMVELAKVFLRRGMAVVMAIVDSPDKDSVSAEALARLAAANPDIAFRHLPVPSRGTERCSTNPVMRAIDVLRAANPALLGFLRALPAVDALVLDMFCTDALDVAAELGVPAYIFFSSALGDLAVMLHLPYYYPTAPSSFKDTPETVLHFPGVPPIRALDMGATMQDRDSDVAKARLSQCARMLEARGILVNSFDWLEARALEALSRGLCTPGRSAPPVHCIGPLVLAGNKGGASERHACLEWLDAQPDRSVVFLSFGSLGRFSMPQLREIARGLENSGQRFLWVVRSPPEHRSNSVEPDLDLEPLLPEGFLERTRERGFAVKNWAPQSEVLRHLSIGAFVTHCGWNSALEGIASGVPMICWPLYAEQKMNKVHMVEELKVGVVMEGYEEELVKAEEVEAKVRLVMAPGSGDGEELRQRLVTAKDMAVEVLKEGGSSHVAFDAFLTDLLKNTRTEKSGFL

>D7S016

MAQIPHIAILPSPGMGHLIPLVEFAKRIFLHHHFSVSLILPTDGPISNAQKIFLNSLPSSMDYHLLPPVNFDDLPEDVKIETRISLTVSRSLTSLRQVLESIIESKKTVALVVDLFGTDAFDVAIDLKISPYIFFPSTAMGLSLFLHLPNLDETVSCEYRDLPDPIQIPGCTPIHGKDLLDPVQDRNDESYKWLLHHAKRYGMAEGIIVNSFKELEGGAIGALQKDEPGKPTVYPVGPLIQMDSGSKVDGSECMTWLDEQPRGSVLYISYGSGGTLSHEQLIEVAAGLEMSEQRFLWVVRCPNDKIANATFFNVQDSTNPLEFLPKGFLERTKGFGLVLPNWAPQARILSHESTGGFLTHCGWNSTLESVVHGVPLIAWPLYAEQKMNAVMLSEDIKVALRPKVNEENGIVGRLEIAKVVKGLMEGEEGKGVRSRMRDLKDAAAKVLSEDGSSTKALAELATKLRKKCQMIDVANH

>Q4R1I9

MGGDAIVLYPYPGLGHLISMVELGKLLLTHHPSFSITILASTAPTTIAATAKLVASSNDQLTNYIKAVSADNPAINFHHLPTISSLPEHIEKLNLPFEYARLQIPNILQVLQTLKSSLKALILDMFCDALFDVTKDLNIPTFYFYTSAGRSLAVLLNIPTFHRTTNSLSDFGDVPISISGMPPIPVSAMPKLLFDRSTNFYKSFLSTSTHMAKSNGIILNTFDLLEERALKALRAGLCLPNQPTPPIFTVGPLISGKSGDNDEHESLKWLNNQPKDSVVFLCFGSMGVFSIKQLEAMALGLEKSGQRFLWVVRNPPIEELPVEEPSLEEILPKGFVERTKDRGLVVRKWAPQVEVLSHDSVGGFVTHCGWNSVLEAVCNGVPMVAWPLYAEQKLGRVFLVEEMKVAVGVKESETGFVSADELEKRVRELMDSESGDEIRGRVSEFSNGGVKAKEEGGSSVASLAKLAQLWKQK

>B2NID2

MEGVILLYSSAEHLNSMLLLATFIAKHHPSIPITILSSADYSAAASVSTLPSITYRRLPPVAIPPDSIKNPVEAFFEIPRLQNPNLRVALEEISQKTRIRAFVIDFFCNSAFEVSTSLSIPTYFYVSTGSAGVCIFLYFPTTDETVATDIGDLRDFLEFPGSPIIHSSDLPQLTFFRRSNVFKHMLDTSKNMQKSSGILTNGFDAMEFRAKEALTNGLCVPNGPTPPVYLVGPLVAGSNAKKDHECLLWLDRQPSKSVVFLCFGRRGLFSGKQLREMAVALERSGYRFLWSVRNPPENRSPAEDPDLDELLPEGFLERTKDIGFVVKSWAPQKEVLSHDAVAGFVTHCGRSSILEALVNGKPMIGWPMYAEQRMNKVFMVDEMKVALPLEEEEDGFVTAVELEKRLRQLMESKTGRDVRHRVAEMKAAATAAMGENGSAVVALRKFIDSVTRD

>Q8LJ11

MDAGDAATTRARKPVVLYPSPGMGHLVSMIELGKVFAARGLAVTVVVVDPPYGNTGATGPFLAGVTAANPAMTFHRLPKVEVPPVASKHHESLTFEVTRLSNPGLRDFLAGASPVVLIIDFFCNAALDVADELGVPAYMFYTSGAEILAFFLYLPVLHAQTTANFGEMGEELVHAPGIPSFPATHSVLPLMERDDPAYAEFLKASADLCRTQGFLVNTFRSLEPRAVETIAAGSCAPPGVSTPPVYCIGPLIKSAEVGENRSEECLAWLDTQPNGSVVFLCFGSIGLFSAEQIKEVAAGLEASGQRFLWVVRSPPSDDPAKKFDKPPEPDLDALLPKGFLERTKGRGLVVKSWAPQRDVLAHAAVGGFVTHCGWNSVLESIVAGVPMLAWPLYAEQRMNRVFLEKEMRLAVAVEGYDDDVGEGTVKAEEVAAKVRWLMESDGGRALLERTLAAMRRAKAALRDGGESEVTLARLVESWREAASA

>Q9FTW1

MAIESAPARNERRGQHVVLLASPGAGHLLPVAELARRIVEYDGFTATIVTHTNFSSAEHSSTFSSLPPSISIAALPEVSVDDLPADARVETRILTVVRRALPHLRDLLRSLLDSPAGVAVFLSDLLSPRALAVAAELGIPRYVFCTSNLMCLTSFLHNPVLDRTTTCEFRDLPGPVLLPGCVPLHGSDLVDPVQDRANPVYRLVIEMGLDYLRADGFLVNTFDAMEHDTAVAFKELSDKGVYPPAYAVGPFVRSPSGKAANDACIRWLDDQPDGSVLYVCLGSGGTLSTEQTAEVAAGLEASGQRFLWVVRYPSDKDKTASYFSVSGDGDGEDSPTNYLPEGFLERTKGTGLAVPMWAPQVEILNHRAVGGFVSHCGWNSTLETVAAGVPMVAWPLYAEQRMNAVMLSSSRAGLALRPSNAREDGVVTRDEVAAVARELITGEKGAAARRKARELREAAAKATRAPGGPSRQAFEAVVGGAWKKAAAAARGGRAGEPDDNGTAVTAQ

>Q6ESW3

METFTADDQRDADAPRPPRVVLLASPGAGHLIPLAELARWLADHHGVAPTLVTFADLEHPDARSAVLSSLPATVATATLPAVPLDDLPADAGLERTLFEVVHRSLPNLRALLRSAASLAALVPDIFCAAALPVAAELGVPGYVFVPTSLAALSLMRRTVELHDGAAAGEQRALPDPLELPGGVSLRNAEVPRGFRDSTTPVYGQLLATGRLYRRAAGFLANSFYELEPAAVEEFKKAAERGTFPPAYPVGPFVRSSSDEAGESACLEWLDLQPAGSVVFVSFGSAGTLSVEQTRELAAGLEMSGHRFLWVVRMPSFNGESFAFGKGAGDEDDHRVHDDPLAWLPDGFLERTSGRGLAVAAWAPQVRVLSHPATAAFVSHCGWNSTLESVAAGVPMIAWPLHAEQTVNAVVLEESVGVAVRPRSWEEDDVIGGAVVTREEIAAAVKEVMEGEKGRGMRRRARELQQAGGRVWSPEGSSRRALEEVAGKWKAAATATAHK

>Q94IF2

MKTTELVFIPAPGMGHLVPTVEVAKQLVDRDEQLSITVLIMTLPLETNIPSYTKSLSSDYSSRITLLQLSQPETSVSMSSFNAINFFEYISSYKDRVKDAVNETFSSSSSVKLKGFVIDMFCTAMIDVANEFGIPSYVFYTSNAAMLGLQLHFQSLSIEYSPKVHNYLDPESEVAISTYINPIPVKCLPGIILDNDKSGTMFVNHARRFRETKGIMVNTFAELESHALKALSDDEKIPPIYPVGPILNLGDGNEDHNQEYDMIMKWLDEQPHSSVVFLCFGSKGSFEEDQVKEIANALERSGNRFLWSLRRPPPKDTLQFPSEFENPEEVLPVGFFQRTKGRGKVIGWAPQLAILSHPAVGGFVSHCGWNSTLESVRSGVPIATWPLYAEQQSNAFQLVKDLGMAVEIKMDYREDFNKTNPPLVKAEEIEDGIRKLMDSENKIRAKVMEMKDKSRAALLEGGSSYVALGHFVETVMKN

>A6XNC2

MKDTIVLYPAFGSGHLMSMVELGKLILTHHPSFSIKILILTPPNQDTNTINVSTSQYISSVSNKFPSINFHYIPSISFTFTLPPHLQTLELSPRSNHHVHHILQSIAKTSNLKAVMLDFLNYSASQVTNNLEIPTYFYYTSGASLLCLFLNFPTFHKNATIPIKDYNMHTPIELPGLPRLSKEDYPDEGKDPSSPSYQVLLQSAKSLRESDGIIVNTFDAIEKKAIKALRNGLCVPDGTTPLLFCIGPVVSTSCEEDKSGCLSWLDSQPGQSVVLLSFGSLGRFSKAQINQIAIGLEKSEQRFLWIVRSDMESEELSLDELLPEGFLERTKEKGMVVRNWAPQGSILRHSSVGGFVTHCGWNSVLEAICEGVPMITWPLYAEQKMNRLILVQEWKVALELNESKDGFVSENELGERVKELMESEKGKEVRETILKMKISAKEARGGGGSSLVDLKKLGDSWREHASWTSVSPNSPFLFA

>B5SX65

MNLASNFMDKTIHIAVVPGVGYGHLVPILHFSKLLIQLHPDIHVTCIIPTLGSPPSSSETILQTLPSNIDYMFLPEVQPSDLPQGLPMEIQIQLTVTNSLPYLHEALKSLALRIPLVALVVDAFAVEALNFAKEFNMLSYIYFCAAASTLAWSFYLPKLDEETTCEYRDLPEPIKVPGCVPLHGRDLLTIVQDRSSQAYKYFLQHVKSLSFADGVLVNSFLEMEMGPINALTEEGSGNPSVYPVGPIIQTVTGSVDDANGLECLSWLDKQQSCSVLYVSFGSGGTLSHEQIVELALGLELSNQKFLWVVRAPSSSSSNAAYLSAQNDVDALQFLPSGFLERTKEEGFVITSWAPQIQILSHSSVGGFLSHCGWSSTLESVVHGVPLITWPMFAEQGMNAVLVTEGLKVGLRPRVNENGIVERVEVAKVIKRLMEGEECEKLHNNMKELKEVASNALKEDGSSTKTISQLTLKWRNLVQKNQI

>I2BH23

MEDVVSTISRKANLRVVMVPSPGRGHLIPFVELSKRLLLRHNFAITILIPDNGSDMIPQRQFLQSLNLPPTISPLYLPPVSLSDLPSDADSITRVPLTVIRSLPAIRDAIINLQHSGEGLCGRVVAVVVDFLGADALQVATQLQIPPYVFYTCSAFHLTLGLNAPQLLHPTHQEDSTKLLKLPGCIPLLGADLPEPYIDKKKDAYKWMVHSHERISSDAVGIIINSFVDLESDIFKALTEERFRTGSGPTVYPIGPLKRLDSDEDLNQFSNESIDCLEWLDKQPESSVLLISFGSGIGARQSKAQFDELAHGLAMSGKRFIWVVKPPGNDVVPWNSSFLPEGFLKKTKGVGLVIPDWVPQIRILSHGSTGGFMSHCGWNSSLESITNGVPVLAWPQHADQKMNAALLVEDAKVALRVDQSSGEDGIVGREEIARYVKAVLDGDEAKLLRKKMRELKVAANNATGNDGSSTKSLDEVANLWKNQNPY

>I2BH18

MNPTPAEAAATLQLVLVPSPGAGHVFPMVELANQLLNRYPALAVTVCIMKMPFKSTSFDFATYKSSHVDRIKFIDLDPPTLDPNTPPSKRFSSFLEGHAPQVKKILSEHVAASNVSPSVVLDMFCTSFMADAKELGVPSYVFYTFSATFLGLMFQLQALYDEGRFNPVQIKDSDTEFVEISSLKTPIPGNLLPSAVVEPDLLLSLITHTRRTKEYASGILINTFQDFESHAIASLNAGQSQSQTPPPIYPVGPIMELKVKDADHSAGPIMEWLDQQPESSVVFLCFGSMGSFDEEQVNEIAAALEKSGCRFIWSLRRPPPKSGGVKFPTDYEDVTEALPAGFLDRTRGVGKVIGWAPQTMILAHPSTGGFVSHCGWNSVLESMWFGVPVATWPMYAEQQLNAVLLVRELEMAEEIRMSYRKESGEVIKAEEIEKGIMGLMSEESGGERRKKTKEMSEKSRKTVENGGASYHSIGRFVGDD

>I2BH16

MAHTDSNLGTLKHTTKRAELVFIPSPGVGHITALAQLAQLLVARDDNLWITILIMHLPHGDANYTNHTTALASTSSALSDRVKFVDLPPNDAAVDPAAKDVVSFFMYSYKSHIRDAVSKLVDQSPFLSGFLVDMFCTTFIDVAVEFGLPSYVFYTSGAGCLNLTLYFQNLRDAQNVPVSDFNNPVADWKIEGFANSIPGKVLPRPVLNPYQCDGFLNFVQNYRNAKGIVINTFPELESATIEHLSKGGNPPVYPVGPILELKRGGGDVKDKGRSSDIMNWLNEQPPSSVVFLCFGSNGCFNEKQVKQIAEALERAGYRFLWSLRRPPPKGTVSFPLDYENPSDVLPEGFLERTTGLGKIIGWAPQAAILAHSAVGGFVSHCGWNSILESLWFGVPIATWPIDGEQQLNAFEMVKEWGLGVDIKMEYSKEFGVDEDDVITVSSDEIEKGLKGLMEDQGGEVRERVRKLSDKCREALAEGGSADIALNGFITDAIRS

>A0A286Q1T2

MEDMQNQPPHVVLLSSPGLGHLIPIIELGKRFLRHHNFKVTILAVTFQASNAEAEVLRSSLCDVIEIPPPDLPAGAGAALATRLCMTMRAAVPAIRAVLSTMQLRPSAFIVDIFGTDSLPVAAELNIRKFVYVTSHAWFLSLLVYLSVLDEIIEGQYVDQNKPLTIPGSSPVRPEDVFDPMQDRNDMQYSECLTLGKAIPRSDGVLVNTWEELQCRDLEALRDQNLMGRVLKVPVYAVGPIMREPDSETGWDSEWVVQWLDKQPRDSVVYVSFGSGGTMSYEQMREIAFGLELSEQRFVWVVRAPTEGVTDAAYLTMGRGTSDDWDGVLPEGFVERTWEVGLVVPQWAPQVTVLRHSSVGAFLSHCGWGSTLESVMSGVPMIAWPLYAEQRMNATLLAEELGVAVRTKELPTKKVVRREEIASMVREVIQLRDGRISAVTKRVKEIKHSAEKALSPGGSSHTALAQVAKIIIGG

>A0A186MUM5

MEKKIHIAVVPSIGFSHLAPILQFSKRLVHLHPHFHVTCLIPSLGSPPLASQTILQNLPPNITYTFLPPVNPNDLPQGIDIVIQIQLTLTHSLPSIHQALNSLTIRIPHVALVVDSLALVALDFAQEFNMLSYVYFPAAATTLSLHFYVPKLDKETSCEYRDLPDPIQIPGCVPLHGKDLYTPAQERSGQTYQSLLQRINRYCSVDGVFVNSFLELETGPIRALTEEGRGYPPVYPVGPIIQAGAKSSDDDDANRLKLKCLAWLDLQQPCSVLYVSFGSGGTLSLEQTLELALGLELSNHKFLWVVRAPSSSASGAYLSAQNDDDPLQFLPSGFLERTKGQGLVIPSWAPQIEILSHCSVGGFLSHCGWNSTLESVVHGVPLITWPLFAEQRMNAVLLCDGLKVGLRPRVGENGLVERVEVAEVVKCLMEREEGGKLHRRMKELKEAASNALKEDGSSTKTLSQLILKWKKLV

>B6EWZ3

MAETPVVTPHIAILPSPGMGHLIPLVEFSKRLIQNHHFSVTLILPTDGPVSNAQKIYLNSLPCSMDYHLLPPVNFDDLPLDTKMETRISLTVTRSLPSLREVFKTLVETKKTVALVVDLFGTDAFDVANDFKVSPYIFYPSTAMALSLFLYLPKLDETVSCEYTDLPDPVQIPGCIPIHGKDLLDPVQDRKNEAYKWVLHHSKRYRMAEGIVANSFKELEGGAIKALQEEEPGKPPVYPVGPLIQMDSGSGSKADRSECLTWLDEQPRGSVLYISFGSGGTLSHEQMIELASGLEMSEQRFLWVIRTPNDKMASATYFNVQDSTNPLDFLPKGFLEKTKGLGLVVPNWAPQAQILGHGSTSGFLTHCGWNSTLESVVHGVPFIAWPLYAEQKMNAVMLSEDIKVALRPKANENGIVGRLEIAKVVKGLMEGEEGKVVRSRMRDLKDAAAKVLSEDGSSTKALAELATKLKKKVSNN

>A6BM07

MKDTIVLYPNLGRGHLVSMVELGKLILTHHPSLSITILILTPPTTPSTTTTTLACDSNAQYIATVTATTPSITFHRVPLAALPFNTPFLPPHLLSLELTRHSTQNIAVALQTLAKASNLKAIVIDFMNFNDPKALTENLNNNVPTYFYYTSGASTLALLLYYPTIHPTLIEKKDTDQPLQIQIPGLSTITADDFPNECKDPLSYACQVFLQIAETMMGGAGIIVNTFEAIEEEAIRALSEDATVPPPLFCVGPVISAPYGEEDKGCLSWLNLQPSQSVVLLCFGSMGRFSRAQLKEIAIGLEKSEQRFLWVVRTELGGADDSAEELSLDELLPEGFLERTKEKGMVVRDWAPQAAILSHDSVGGFVTHCGWNSVLEAVCEGVPMVAWPLYAEQKMNRMVMVKEMKVALAVNENKDGFVSSTELGDRVRELMESDKGKEIRQRIFKMKMSAAEAMAEGGTSRASLDKLAKLWKQS

>F8WKX1

MENTLVLYPAPGIGHMISMLELAKLILRHYSNKFSRIHILINTGFRDMKSTYLDHISSTNPSIVVHQFPFIQADLSSSLSPPAIGFKFIRKNAPNVHHALQEISKTSSIRALIIDFFCTSAMPYSNNLGIPVYYFFTSGAAAVALFLYFPTIHKQTSESFKDLVQTKFDVPGLPPIPATQMPEPVLDRDDPAYDDILYYSVHLPKSSGIIVNTFDELEPIALKAITDGLCVPDAPTPPLYNIGPLIADADSRPAIDGDKGIDLDQSDCFSWLDRQPDQCVVFLCFGSRGTFSVEQIKEIAKGLERSGKRFLWVVKKPLRNNKSKQVEGSGGFEIDSILPERFLEKTKGIGLVVKSWIPQLQVLRHPAVGGFVTHCGWNSTLEAVVAGVPLVAWPLHAEQHVNMAALVQDMKMAIPVEQGDDGIVRGEEVEKRVRELMDSERGRELRKLSQKTRDIAAESGVHLGSSSTALASLIHVVFGN

>F8WKX0

MKNQELVFIPSAVMSHLVSTVELAKLLIDRNEHLSITVLIMKLPYEKNVGNYTYPQTEASDSRIRFLELKKDESASQTVSPILFIYQFVEDHKNSARDVLTEISNSASSDLVGVVVDMFCSSMIDVANEFGVPSYVFYTSGAAMLGLMLHLQSLRDDFGEDVTNYENSKVELAVPTYINPVPVKVLPSRLFDMEGGGNMFLNLTKRFRETKGIVINSFFELESHAIQALSNDKTIPPVYPVGPILDLKESNGQNQETEMITKWLDIQPDSSVVFLCFGSRGCFDGGQVKEIACALESSGYRFLWSLRRPPPKGKFESPGDYENLEEALPEGFLQRTAEVGKVIGWAPQAAILSHPAVGCFVSHCGWNSTLESVWFGVPMATWPLYAEQQVNAFLLLKDLGMAVDIKMDFKSTSFEPSTEIVAADLIEKAIKHLMDPENEIRKKVKEKKEKSRLSLSEGGPSSASLGRFLDALIDNIP

>D2KY82

METIVLYPSPGMGHLISMVELGKFILKHHPSFTIAILIVPPSFNTGSTASYIDRVSAATNSITFHHLPTISLELDSFSSMEALIFEAIRLSNPHVHHALQHISLTTTITALIIDFFCTPAISISTKLGIPTYYFFTSGISSLAFFLYLPVIHRNTVKSFKDLNSLVDIPGLPPIPSSDVAKPILDRASTEYACFLDFSLHLPKSAGVIVNSFNSLEPKTLKAISEGSCNPDGATPPVFCVGPLLATEDQQSGTDGVHECLKWLDLQPIQSVVFLCFGSLGLFSDKQLKEIAIGLERSEQRFLWVVRSPPSEDKSKRFLAPPEPDLDSLLPIGFLDRTKDLGFVVKSWAPQVEVLNHKSIGGFVTHCGWNSVLEAVCAGVPMVAWPLYAEQKFNRVILVEDLKLALRINESEDGFVTAEEVESRVRELMDSDEGESLRKLAKEKEAEAKAAISEGGSSIVDLAKLVESWKLH

>A0A0A1H7N8

MEEAIVFYSAPGMGHIVSMAELANLIRRHLLLHYPNRRVSFTVIYGSQPFEPPTTTTAISQIANSSPSISFLHLPQHPINTFPARSPAAMAFESIRKSAPDFAESLRRISDAGTKIRSLVIDLFCGTALPIAAEQGIPVYYFFTSGSAALAAYLHMPTIHQRVGGRVFKDSPDLLIDIPGLPSIPADEMPEPLLDAKDPAYPEMVYFCECLQKSKGILVNTFDELEPVVTREAIDSGACVPDGPTPAVYNIGPLIEGSKDGDTSHETLSWLNTQPSGSVVFLCFGSRGRFTAAQTKAIAEGLERSGQRFIWVVRNPPDDNSSEIDLGKLLPEGFLRRTKDIGIVVKGWAPQVAVLSHDSVGGFVTHCGWNSVLEAVVAGKPMVAWPLYAEQHLNRAVLVKTMGMAVPVEQREGDRFVSGDELASRLIELMNSEKGKEMKEKSRLMRDKALAARNEGGSSMADLQKVVNEWMK

>A0A0A1H7P3

MSSSSAPVPPTTTPELAAGQPPHVVIFPSPGMGHLIPLTEFAKRLIPRFTFTFAVPTAGEPSSAQRSFLSALPAGIDSVFLPSVSMSDVPSDARIETLMSIMVSRSLPSLRDLLLSRSPVAVLLVDLFGTDAFDVAFEVGITPYIFFPSTAMGLSLFLHLEKLDESVSCEYADLTEPVKIPGCVPVHGRDLLDPVQDRKNDAYKWVLHHCKRYKLAKGIIVNSFEAVERGAIRALVQAEPAVYPVGPLVQTCSPEQPTECIKWLDRQPRGSVLFVNFGSGGTLSTEQQKELALGLARSEQRFLWVVRSPNDAVANATYFSVDGESDPLQLLPSGFLDETAGRGLVVPKWAPQIEVLSHGSTGGFLTHCGWNSILESVVHGVPLITWPLYAEQKMNAVMLTEDLKVGLRPVAGKDGIIRGDEIARVVRELMEGEEGKRIKDRMVELKVAAKVALSEDGSSTKALIEIAKQWETKV

>A0A0A1H7L4

MERPHIFMLPSPGMGHFIPLIELAKILASHHRVSATILVPTTGSGHLTPAQKPFLDSLPNGINYLLLPPVELDDVVDDARFEVRVLLLMSRSIPFVRDAMAGCRISAFVADPFGIDAFEIAKEFGIPSYLYFPASATSLVFFLHLPDLDGSVFGEFGDLPSPVLLPGCAPLHGKDLLDSVQDRKNEAYQWILHLSKKARVLPDGIMLNSFAMLEPGPIRALLQEKLPGPTAIYPIGPVIQSGSVKEVDRVEREECLTWLDDQPDGSVLFISFGSGGTLSYDQHTELALGLEKCDYIRFIWVIRIPNCKSSNRPLLSPKCDHNPLEFLPRGFVERTKARGMVMNWWAPQIDILSHRSTGGFLTHCGWNSTLEAIVHGVPLIGWPLYAEQKMNAMMLHETLEIALMPQADPVSGLTDNEEIARVVKDLMEGEDGKRVRHRIKDLSEATKKFNSEDGDSTKLLAQVILKWGDYNN

>A0A0A1H9X0

MKKAELVFIPSPGAGHTVSAAEFANRLIARDARISVTILTMSSPFTTAASSTSTYPNPEPSVDVPRYISLPQVQLPPIKILQDSIEEYISTFAASHSHFVRNELISLQKSPEIPVYLVVDMFCTAFIDVANELGIPSYVFFTSGSAFLGLRLNLPDLYERTGIAKFIESEPEFEIPSFRNPVPASGFPGFAFNKSGYTSFMNHARKFQETKGIMVNTFAELEPYAIKSLTEISPNKVPVVYTIGPVLNLKGAAHKPCDQDIRARIMNWLDIQPESSVVYLCFGSVGSFEEEQVKEIAKGLNQSGQRFLWSLRKRITEGAIRPTDYTDEELKSVLPDDFRTFVAEGRGMVCGWAPQVEILAHKAIGSFVTHCGWNSTLESVWFGKPIVAWPLYAEQRVNAFELVKELGLAAAWLGYGTKPGDVVTANEVESAVRSVMDKDNPVRSKVKEISEVSRLTLLDGGSSFNSIGSFIKNIIPEEL

>A7M6I2

MSSSTELVIVPAPGMGHLVSTVQLAKVILKKYDFISISIFIINLPMHSDKISSYVDSQSRDNPYPTRLLFTTLPPVTITSDPTSLGFFTDFIKLHKPLVKRAVEERVELGSPKPAGFVLDMFCTTMVDVANELGIPSYLFLTCGVNFLNFVYYVESLADEHGLGAREVSAKLSDPEFESVVSGFRNPITSKIIPGIFKGEFGSGMILNLAKEFKKMKGILVNSYVELESFEIQALQNSDDKKIPPIYPVGPILDLNRESGSDKEENKSIIEWLNSQPDSSIVFLCFGSMGSFDAEQVKEIANGLEKSGVRFLWALRKPPSPDQRGPPSDNGTFLEALPEGFIDRTVNRGKIIGWAPQVDVLAHPAIGGFVSHCGWNSTLESLWFGVPIGAWPMYSEQNLNALVLVEQKLAVEIRMDYVMDWLSKKGNFIVSSMEIEEGLKKLMNMDENMRRNVKDMGEKGRKALEKGGSSCHWLDSFMKDVLTNVA

>A0A1P8C3B1

MGPHQKTMTNSELVFIPSPGAGHLPPTVELAKLLLRRHHRLSITIIIMKAPFGGGAYDTAKLDSTPRLRCVEIPSDDSTAALISPTAFLTAFIDHHKPHVRNIVRESIADPGGTVRLAGFVVDMFCVDMVDVANEFGAPTYAYFTSGAAMLGLMFHLQAKRDDDNLDVTELDKNSRSVISVPSYINPVPVNVLLDVLFDKNGAKMFLDLAKRFRETKGIIVNSFRELESHAIEVLSDDPDIPPVFPVGPILNLNSTTDDGKVDDIMTWLDEQPERSVVFLCFGSMGTFPEEQIREIASAIESGGHRFLWSLRRPSSKEKMESPKEYEDPGEVLPEGFLERTSGVGRVIGWAPQLMVLSHRSVGSFVSHCGWNSTLESIWCGVPIAAWPMYAEQQTNAFQLVSEAGIAAEVRMDYRTNMKPGGEQMIVTAEEIERGIRRVMSDGEIKRKAEEMKEKSRLAVSEGGSSYDSIGDFIHHVMNE

>B9VNV0

MEDAIVLYSSAEHLNSMLVPAKFISKHHPSISVIIISTAAESAAASVASVPSITYHRLPSAPLPPDLTTSIIELFFEIPRFHNPFLHEALLEISQKSNLRAFLIDFFCNSTFEVSTSLNIPTYFYLSGGACGLCALLYFPTIDEAVSPRDIGELNDFLEIPGCPPVHSLDFPKAMWFRRSNTYKHFLDTAGNMRRASGIVTNSFDAIEFRAKEALSNSLCTPGLATPPVYVIGPLVAETNRKNGGEEHECLKWLDSQPIKSVIFLCFGRRGLFSAAQLKEMAIGLENSGHRFLWSVRSPPGPAAAKDPDLDALLPEGFMERTKDRGFVIKTWAPQKEVLSHEAVGGFVTHCGRSSVLEAVSFGVPMIGWPMYAEQRMQRVFMVEEMKVALPLAEEADGFVTAGELEKRVRELMGLPAGKAVTQRVAELRTAAEAAVRKGGSSVVALGKFIETVTRR

>O82383

MRNVELIFIPTPTVGHLVPFLEFARRLIEQDDRIRITILLMKLQGQSHLDTYVKSIASSQPFVRFIDVPELEEKPTLGSTQSVEAYVYDVIERNIPLVRNIVMDILTSLALDGVKVKGLVVDFFCLPMIDVAKDISLPFYVFLTTNSGFLAMMQYLADRHSRDTSVFVRNSEEMLSIPGFVNPVPANVLPSALFVEDGYDAYVKLAILFTKANGILVNSSFDIEPYSVNHFLQEQNYPSVYAVGPIFDLKAQPHPEQDLTRRDELMKWLDDQPEASVVFLCFGSMARLRGSLVKEIAHGLELCQYRFLWSLRKEEVTKDDLPEGFLDRVDGRGMICGWSPQVEILAHKAVGGFVSHCGWNSIVESLWFGVPIVTWPMYAEQQLNAFLMVKELKLAVELKLDYRVHSDEIVNANEIETAIRYVMDTDNNVVRKRVMDISQMIQRATKNGGSSFAAIEKFIYDVIGIKP

>Q9LML6

MVKETELIFIPVPSTGHILVHIEFAKRLINLDHRIHTITILNLSSPSSPHASVFARSLIASQPKIRLHDLPPIQDPPPFDLYQRAPEAYIVKLIKKNTPLIKDAVSSIVASRRGGSDSVQVAGLVLDLFCNSLVKDVGNELNLPSYIYLTCNARYLGMMKYIPDRHRKIASEFDLSSGDEELPVPGFINAIPTKFMPPGLFNKEAYEAYVELAPRFADAKGILVNSFTELEPHPFDYFSHLEKFPPVYPVGPILSLKDRASPNEEAVDRDQIVGWLDDQPESSVVFLCFGSRGSVDEPQVKEIARALELVGCRFLWSIRTSGDVETNPNDVLPEGFMGRVAGRGLVCGWAPQVEVLAHKAIGGFVSHCGWNSTLESLWFGVPVATWPMYAEQQLNAFTLVKELGLAVDLRMDYVSSRGGLVTCDEIARAVRSLMDGGDEKRKKVKEMADAARKALMDGGSSSLATARFIAELFEDGSSC

>O82382

MAKQQEAELIFIPFPIPGHILATIELAKRLISHQPSRIHTITILHWSLPFLPQSDTIAFLKSLIETESRIRLITLPDVQNPPPMELFVKASESYILEYVKKMVPLVRNALSTLLSSRDESDSVHVAGLVLDFFCVPLIDVGNEFNLPSYIFLTCSASFLGMMKYLLERNRETKPELNRSSDEETISVPGFVNSVPVKVLPPGLFTTESYEAWVEMAERFPEAKGILVNSFESLERNAFDYFDRRPDNYPPVYPIGPILCSNDRPNLDLSERDRILKWLDDQPESSVVFLCFGSLKSLAASQIKEIAQALELVGIRFLWSIRTDPKEYASPNEILPDGFMNRVMGLGLVCGWAPQVEILAHKAIGGFVSHCGWNSILESLRFGVPIATWPMYAEQQLNAFTIVKELGLALEMRLDYVSEYGEIVKADEIAGAVRSLMDGEDVPRRKLKEIAEAGKEAVMDGGSSFVAVKRFIDGL

>O82381

MGKQEDAELVIIPFPFSGHILATIELAKRLISQDNPRIHTITILYWGLPFIPQADTIAFLRSLVKNEPRIRLVTLPEVQDPPPMELFVEFAESYILEYVKKMVPIIREALSTLLSSRDESGSVRVAGLVLDFFCVPMIDVGNEFNLPSYIFLTCSAGFLGMMKYLPERHREIKSEFNRSFNEELNLIPGYVNSVPTKVLPSGLFMKETYEPWVELAERFPEAKGILVNSYTALEPNGFKYFDRCPDNYPTIYPIGPILCSNDRPNLDSSERDRIITWLDDQPESSVVFLCFGSLKNLSATQINEIAQALEIVDCKFIWSFRTNPKEYASPYEALPHGFMDRVMDQGIVCGWAPQVEILAHKAVGGFVSHCGWNSILESLGFGVPIATWPMYAEQQLNAFTMVKELGLALEMRLDYVSEDGDIVKADEIAGTVRSLMDGVDVPKSKVKEIAEAGKEAVDGGSSFLAVKRFIGDLIDGVSISK

>Q9LSY9

MKVELVFIPSPGVGHIRATTALAKLLVASDNRLSVTLIVIPSRVSDDASSSVYTNSEDRLRYILLPARDQTTDLVSYIDSQKPQVRAVVSKVAGDVSTRSDSRLAGIVVDMFCTSMIDIADEFNLSAYIFYTSNASYLGLQFHVQSLYDEKELDVSEFKDTEMKFDVPTLTQPFPAKCLPSVMLNKKWFPYVLGRARSFRATKGILVNSVADMEPQALSFFSGGNGNTNIPPVYAVGPIMDLESSGDEEKRKEILHWLKEQPTKSVVFLCFGSMGGFSEEQAREIAVALERSGHRFLWSLRRASPVGNKSNPPPGEFTNLEEILPKGFLDRTVEIGKIISWAPQVDVLNSPAIGAFVTHCGWNSILESLWFGVPMAAWPIYAEQQFNAFHMVDELGLAAEVKKEYRRDFLVEEPEIVTADEIERGIKCAMEQDSKMRKRVMEMKDKLHVALVDGGSSNCALKKFVQDVVDNVP

>Q33DV3

MGEEYKKTHTIVFHTSEEHLNSSIALAKFITKHHSSISITIISTAPAESSEVAKIINNPSITYRGLTAVALPENLTSNINKNPVELFFEIPRLQNANLREALLDISRKSDIKALIIDFFCNAAFEVSTSMNIPTYFDVSGGAFLLCTFLHHPTLHQTVRGDIADLNDSVEMPGFPLIHSSDLPMSLFYRKTNVYKHFLDTSLNMRKSSGILVNTFVALEFRAKEALSNGLYGPTPPLYLLSHTIAEPHDTKVLVNQHECLSWLDLQPSKSVIFLCFGRRGAFSAQQLKEIAIGLEKSGCRFLWLARISPEMDLNALLPEGFLSRTKGVGFVTNTWVPQKEVLSHDAVGGFVTHCGWSSVLEALSFGVPMIGWPLYAEQRINRVFMVEEIKVALPLDEEDGFVTAMELEKRVRELMESVKGKEVKRRVAELKISTKAAVSKGGSSLASLEKFINSVTR

>A0A4Y5RXX8 BLAST

MEKSNPNSTSKPHVFLLASPGMGHLIPFLELSKRLVTLNTLQVTLFIVSNEATKARSHLMESSNNFHPDLELVDLTPANLSELLSTDATVFKRIFLITQAAIKDLESRISSMSTPPAALIVDVFSMDAFPVADRFGIKKYVFVTLNAWFLALTTYVRTLDREIEGEYVDLPEPIAIPGCKPLRPEDVFDPMLSRSSDGYRPYLGMSERLTKADGLLLNTWEALEPVSLKALRENEKLNQIMTPPLYPVGPVARTTVQEVVGNECLDWLSKQPTESVLYVALGSGGIISYKQMTELAWGLEMSRQRFIWVVRLPTMEKDGACRFFSDVNVKGPLEYLPEGFLDRNKELGMVLPNWGPQDAILAHPSTGGFLSHCGWNSSLESIVNGVPVIAWPLYAEQKMNATLLTEELGVAVRPEVLPTKAVVSRDEIEKMVRRVIESKEGKMKRNRARSVQSDALKAIEKGGSSYNTLIEVAKEFEKNHKVL

>Q5IFH7 BLAST

MSMSDINKNSELIFIPAPGIGHLASALEFAKLLTNHDKNLYITVFCIKFPGMPFADSYIKSVLASQPQIQLIDLPEVEPPPQELLKSPEFYILTFLESLIPHVKATIKTILSNKVVGLVLDFFCVSMIDVGNEFGIPSYLFLTSNVGFLSLMLSLKNRQIEEVFDDSDRDHQLLNIPGISNQVPSNVLPDACFNKDGGYIAYYKLAERFRDTKGIIVNTFSDLEQSSIDALYDHDEKIPPIYAVGPLLDLKGQPNPKLDQAQHDLILKWLDEQPDKSVVFLCFGSMGVSFGPSQIREIALGLKHSGVRFLWSNSAEKKVFPEGFLEWMELEGKGMICGWAPQVEVLAHKAIGGFVSHCGWNSILESMWFGVPILTWPIYAEQQLNAFRLVKEWGVGLGLRVDYRKGSDVVAAEEIEKGLKDLMDKDSIVHKKVQEMKEMSRNAVVDGGSSLISVGKLIDDITG

>A6XNC5 BLAST

MGNFANRKPHVVMIPYPVQGHINPLFKLAKLLHLRGFHITFVNTEYNHKRLLKSRGPKAFDGFTDFNFESIPDGLTPMEGDGDVSQDVPTLCQSVRKNFLKPYCELLTRLNHSTNVPPVTCLVSDCCMSFTIQAAEEFELPNVLYFSSSACSLLNVMHFRSFVERGIIPFKDESYLTNGCLETKVDWIPGLKNFRLKDIVDFIRTTNPNDIMLEFFIEVADRVNKDTTILLNTFNELESDVINALSSTIPSIYPIGPLPSLLKQTPQIHQLDSLDSNLWKEDTECLDWLESKEPGSVVYVNFGSTTVMTPEQLLEFAWGLANCKKSFLWIIRPDLVIGGSVIFSSEFTNEIADRGLIASWCPQDKVLNHPSIGGFLTHCGWNSTTESICAGVPMLCWPFFADQPTDCRFICNEWEIGMEIDTNVKREELAKLINEVIAGDKGKKMKQKAMELKKKAEENTRPGGCSYMNLNKVIKDVLLKQN

>O22822 BLAST

MEHKRGHVLAVPYPAQGHITPFRQFCKRLHFKGLKTTLALTTFVFNSINPDLSGPISIATISDGYDHGGFETADSIDDYLKDFKTSGSKTIADIIQKHQTSDNPITCIVYDAFLPWALDVAREFGLVATPFFTQPCAVNYVYYLSYINNGSLQLPIEELPFLELQDLPSFFSVSGSYPAYFEMVLQQFINFEKADFVLVNSFQELELHENELWSKACPVLTIGPTIPSIYLDQRIKSDTGYDLNLFESKDDSFCINWLDTRPQGSVVYVAFGSMAQLTNVQMEELASAVSNFSFLWVVRSSEEEKLPSGFLETVNKEKSLVLKWSPQLQVLSNKAIGCFLTHCGWNSTMEALTFGVPMVAMPQWTDQPMNAKYIQDVWKAGVRVKTEKESGIAKREEIEFSIKEVMEGERSKEMKKNVKKWRDLAVKSLNEGGSTDTNIDTFVSRVQSK

>P51094 BLAST

MSQTTTNPHVAVLAFPFSTHAAPLLAVVRRLAAAAPHAVFSFFSTSQSNASIFHDSMHTMQCNIKSYDISDGVPEGYVFAGRPQEDIELFTRAAPESFRQGMVMAVAETGRPVSCLVADAFIWFAADMAAEMGVAWLPFWTAGPNSLSTHVYIDEIREKIGVSGIQGREDELLNFIPGMSKVRFRDLQEGIVFGNLNSLFSRMLHRMGQVLPKATAVFINSFEELDDSLTNDLKSKLKTYLNIGPFNLITPPPVVPNTTGCLQWLKERKPTSVVYISFGTVTTPPPAEVVALSEALEASRVPFIWSLRDKARVHLPEGFLEKTRGYGMVVPWAPQAEVLAHEAVGAFVTHCGWNSLWESVAGGVPLICRPFFGDQRLNGRMVEDVLEIGVRIEGGVFTKSGLMSCFDQILSQEKGKKLRENLRALRETADRAVGPKGSSTENFITLVDLVSKPKDV

>Q6VAB4 BLAST

MENKTETTVRRRRRIILFPVPFQGAINPILQLANVLYSKGFSITIFHTNFNKPKTSNYPHFTFRFILDNDPQDERISNLPTHGPLAGMRIPIINEHGADELRRELELLMLASEEDEEVSCLITDALWYFAQSVADSLNLRRLVLMTSSLFNFHAHVSLPQFDELGYLDPDDKTRLEEQASGFPMLKVKDIKSAYSNWQILKEILGKMIKQTKASSGVIWNSFKELEESELETVIREIPAPSFLIPLPKHLTASSSSLLDHDRTVFQWLDQQPPSSVLYVSFGSTSEVDEKDFLEIARGLVDSKQSFLWVVRPGFVKGSTWVEPLPDGFLGERGRIVKWVPQQEVLAHGAIGAFWTHSGWNSTLESVCEGVPMIFSDFGLDQPLNARYMSDVLKVGVYLENGWERGEIANAIRRVMVDEEGEYIRQNARVLKQKADVSLMKGGSSYESLESLVSYISSLLEHHHHHH

>Q7XT97 BLAST

GMGSMSTPAASANGGQVLLLPFPAAQGHTNPMLQFGRRLAYHGLRPTLVTTRYVLSTTPPPGDPFRVAAISDGFDDASGMAALPDPGEYLRTLEAHGARTLAELLLSEARAGRPARVLVYDPHLPWARRVARAAGVATAAFLSQPCAVDLIYGEVCARRLALPVTPTDARGLYARGVLGVELGPDDVPPFVAAPELTPAFCEASIEQFAGLEDDDDVLVNSFSDLEPKEAAYMESTWRAKTIGPSLPSFYLDDGRLRSNTAYGFNLFRSTVPCMEWLDKQPPRSVVLVSYGTVSTFDVAKLEELGNGLCNSGKPFLWVVRSNEEHKLSVQLRKKCEKRGLIVPFCPQLEVLAHKATGCFLSHCGWNSTLEAIVNGVPLVAMPHWADQPTISKYVESLWGMGVRVQLDKSGILQREEVERCIREVMDGDRKEDYRRNATRLMKKAKESMQEGGSSDKNIAEFAAKYSN

>A6XNC6 BLAST

MSTFKNEMNGNNLLHVAVLAFPFGTHAAPLLSLVKKIATEAPKVTFSFFCTTTTNDTLFSRSNEFLPNIKYYNVHDGLPKGYVSSGNPREPIFLFIKAMQENFKHVIDEAVAETGKNITCLVTDAFFWFGADLAEEMHAKWVPLWTAGPHSLLTHVYTDLIREKTGSKEVHDVKSIDVLPGFPELKASDLPEGVIKDIDVPFATMLHKMGLELPRANAVAINSFATIHPLIENELNSKFKLLLNVGPFNLTTPQRKVSDEHGCLEWLDQHENSSVVYISFGSVVTPPPHELTALAESLEECGFPFIWSFRGDPKEKLPKGFLERTKTKGKIVAWAPQVEILKHSSVGVFLTHSGWNSVLECIVGGVPMISRPFFGDQGLNTILTESVLEIGVGVDNGVLTKESIKKALELTMSSEKGGIMRQKIVKLKESAFKAVEQNGTSAMDFTTLIQIVTS

>A4F1R4 BLAST

MKNKQHVAIFPFPFGSHLPPLLNLVLKLAHIAPNTSFSFIGTHSSNAFLFTKRHIPNNIRVFTISDGIPEGHVPANNPIEKLDLFLSTGPDNLRKGIELAVAETKQSVTCIIADAFVTSSLLVAQTLNVPWIAFWPNVSCSLSLYFNIDLIRDKCSKDAKNATLDFLPGLSKLRVEDVPQDMLDVGEKETLFSRTLNSLGVVLPQAKAVVVNFFAELDPPLFVKYMRSKLQSLLYVVPLPCPQLLLPEIDSNGCLSWLDSKSSRSVAYVCFGTVVSPPPQEVVAVAEALEESGFPFVWALKESLLSILPKGFVERTSTRGKVVSWVPQSHVLSHGSVGVFVTHCGANSVMESVSNGVPMICRPFFGDQGIAARVIQDIWEVGVIVEGKVFTKNGFVKSLNLILVQEDGKKIRDNALKVKQIVQDAVGPHGQAAEDFNTLVEVISSS

>O81498 indigo

MHITKPHAAMFSSPGMGHVIPVIELGKRLSANNGFHVTVFVLETDAASVQSKLLNSTGVDIVNLPSPDISGLVDPNAHVVTKIGVIMREAVPTLRSKIVAMHQNPTALIIDLFGTDALCLAAELNMLTYVFIASNARYLGVSIYYPTLDEVIKEEHTVQRKPLTIPGCEPVRFEDIMDAYLVPDEPVYHDLVRHCLAYPKADGILVNTWEEMEPKSLKSLQDPKLLGRVARVPVYPVGPLCRPIQSSTTDHPVFDWLNKQPNESVLYISFGSGGSLTAQQLTELAWGLEESQQRFIWVVRPPVDGSSCSDYFSAKGGVTKDNTPEYLPEGFVTRTCDRGFMIPSWAPQAEILAHQAVGGFLTHCGWSSTLESVLCGVPMIAWPLFAEQNMNAALLSDELGISVRVDDPKEAISRSKIEAMVRKVMAEDEGEEMRRKVKKLRDTAEMSLSIHGGGSAHESLCRVTKECQRFLECVGDLSRGA

>Q9M156 indigo

MEESKTPHVAIIPSPGMGHLIPLVEFAKRLVHLHGLTVTFVIAGEGPPSKAQRTVLDSLPSSISSVFLPPVDLTDLSSSTRIESRISLTVTRSNPELRKVFDSFVEGGRLPTALVVDLFGTDAFDVAVEFHVPPYIFYPTTANVLSFFLHLPKLDETVSCEFRELTEPLMLPGCVPVAGKDFLDPAQDRKDDAYKWLLHNTKRYKEAEGILVNTFFELEPNAIKALQEPGLDKPPVYPVGPLVNIGKQEAKQTEESECLKWLDNQPLGSVLYVSFGSGGTLTCEQLNELALGLADSEQRFLWVIRSPSGIANSSYFDSHSQTDPLTFLPPGFLERTKKRGFVIPFWAPQAQVLAHPSTGGFLTHCGWNSTLESVVSGIPLIAWPLYAEQKMNAVLLSEDIRAALRPRAGDDGLVRREEVARVVKGLMEGEEGKGVRNKMKELKEAACRVLKDDGTSTKALSLVALKWKAHKKELEQNGNH

>A0A024AX02 indigo

MAETAIVTKSENPHIVILPSPGMGHLIPLVEFSKRLISQHQFSVTLILPTDGPISNSQKSFLNSLPSCMDYHLLPPVNFDDLPLDVKIETRISLTVTRSLSSLREVFKTLVDSKKVVAFVVDLFGTDAFDVAIDFNVSPYIFFPSTAMALSLFLYLPKLDATVSCEYRDLPDPIQIPGCIPIHGKDLLDPVQDRKNEAYRWLLHHSKRYRMAEGVVSNSFKELEGGPIKALQEEEPGKPPVYPVGPLIQMDSGSKVDGSGCLTWLDEQPRGSVLYVSYGSGGTLSHEQLIEVASGLEMSEQRFLWVIRCPNDTVANATYFNVQDSTNPLDFLPKGFLERTKGLGLVVPNWAPQAQILSHGSTGGFLTHCGWNSTLESVVHGVPLIAWPLYAEQKMNAVMLTEDIKVALRPKANENGLVGRLEIAKVVKGLMEGEEGKGVRTRMRDLKDAAAKVLSQDGSSTKALAELATKLKNKVLIN

>A0A2L2R220 indigo

MESPAAPPTTAPPPHVIIVPSAGMGHLIPLAEFAKRLLPRFTFTFAVPTSGPPSSSQRDFLSSLPASIDTSFLPEVDLSDAPSDAQIETLMSLMVVRSLPSLRDLIASYSASGRRVAALVVDLFATDAIDVALELGIRPFIFFPSTAMTLSFFLHLEKLDETVSCEFAELSDPVQIPGCIPVHGKDLIDPVQDRKNDAYKWLLHHSKRYKLAEGVIVNSFEGLEGGPIRELLHPEPGKPRVYPVGPLIQAGSCEKGAAARPECLKWLDQQPRGSVLFVNFGSGGVLSTEQQNELAGVLAHSQQRFLWVVRPPNDGIANATYFSVDGEIDPLKLLPEGFLEQTAGRGLVLPMWAPQIDVLSHESTGGFLTHCGWNSTLESVFHGVPLITWPLYAEQKMNAVMLTEGLRVGLRPSVGKDGIIRGAEIARVIGELMEGEEGKRIRSKMQELKRAASAVLSKDGSSTRALEEVAKIWESKV

>Q9LVR1 indigo

MHITKPHAAMFSSPGMGHVIPVIELGKRLSANNGFHVTVFVLETDAASAQSKFLNSTGVDIVKLPSPDIYGLVDPDDHVVTKIGVIMRAAVPALRSKIAAMHQKPTALIVDLFGTDALCLAKEFNMLSYVFIPTNARFLGVSIYYPNLDKDIKEEHTVQRNPLAIPGCEPVRFEDTLDAYLVPDEPVYRDFVRHGLAYPKADGILVNTWEEMEPKSLKSLLNPKLLGRVARVPVYPIGPLCRPIQSSETDHPVLDWLNEQPNESVLYISFGSGGCLSAKQLTELAWGLEQSQQRFVWVVRPPVDGSCCSEYVSANGGGTEDNTPEYLPEGFVSRTSDRGFVVPSWAPQAEILSHRAVGGFLTHCGWSSTLESVVGGVPMIAWPLFAEQNMNAALLSDELGIAVRLDDPKEDISRWKIEALVRKVMTEKEGEAMRRKVKKLRDSAEMSLSIDGGGLAHESLCRVTKECQRFLERVVDLSRGA
